# Maternal care modulates chloride cotransporter development during inhibitory circuit maturation in the piriform cortex

**DOI:** 10.1101/2025.11.05.686805

**Authors:** Christell Becerra-Flores, Enver M. Oruro, Carlos Medina-Saldivar, Grace E. Pardo, Luis F. Pacheco-Otálora

**Author notes:** **Corresponding author** Grace V. E. Pardo, PhD. Laboratorio de Investigación en Neurociencia, Instituto Científico, Universidad Andina del Cusco, Av. Prolongación de La Cultura s/n, Campus Qollana, San Jerónimo, Cusco 08400, Peru. These authors share senior authorship (Grace E. Pardo, Luis F. Pacheco-Otálora).

## Abstract

Early postnatal development is a sensitive period for inhibitory circuit maturation, marked by a shift in GABAergic signaling from depolarizing to hyperpolarizing actions. This transition depends on chloride homeostasis, which is regulated by the potassium-chloride cotransporter 2 (KCC2) and the sodium-potassium-chloride cotransporter 1 (NKCC1). While these cotransporters are known to drive inhibitory development, little is understood about how early caregiving experience influences their trajectories in brain regions critical for attachment learning. Here, we examined the developmental profiles of KCC2 and NKCC1 in the piriform cortex of male and female rats from postnatal (P) day 5 to 22 and assessed their sensitivity to altered maternal care using the limited bedding and nesting (LBN) paradigm from P2 to P9. Both transporters progressively increased protein levels toward adult-like levels, with sex-specific regulation observed at the mRNA level. At P15, LBN reduced KCC2 and NKCC1 protein levels in a region- and sex- dependent manner. To evaluate the functional consequences of these changes at the neural level, we implemented a Hodgkin-Huxley computational model with dynamic ion concentration parameters, which were parameterized using our experimental data. Simulation revealed that cotransporter profiles induced by LBN altered the chloride equilibrium, changing the impact of GABAergic input on neuronal excitability. This integrative approach offers a mechanistic insight into how early caregiving experiences affect chloride transporter regulation, with consequences for synaptic signaling and neuronal activity, ultimately contributing to the development of inhibitory circuits.

## 1. Introduction

Early brain development involves highly dynamic processes of circuit formation, refinement, and plasticity. The coordinated modulation of excitatory and inhibitory neurotransmission is essential for establishing functional neural networks that support cognition, sensory processing, and adaptive behavior throughout life. Among the various molecular systems that regulate early circuit maturation, γ-aminobutyric acid (GABA) signaling plays a particularly prominent role in shaping synaptic organization and network excitability during sensitive periods of postnatal development (Ben-Ari, 2002; Kaila *et al*., 2014).

While GABA serves as the principal inhibitory neurotransmitter in the mature brain, its early postnatal function is predominantly depolarizing. This developmental polarity shift depends on changes in neuronal chloride gradients, which are regulated by cation-chloride cotransporters, primarily the sodium-potassium-chloride importer NKCC1 and the potassium-chloride exporter KCC2 (Payne, 1997; Payne et al., 1996; Rivera et al., 1999). In immature neurons, high intracellular chloride levels maintained by NKCC1 result in depolarizing GABAergic currents (Ben-Ari, 2002; Kaila *et al*., 2014). With the progression of development, the upregulation of KCC2 lowers intracellular chloride concentration, enabling the emergence of hyperpolarizing GABAergic signaling (Ben-Ari, 2002; Payne *et al*., 2003; Kaila *et al*., 2014). The timing and regulation of this GABA switch are closely linked to circuit formation, sensory experience, and the opening of sensitive periods for plasticity (Hensch *et al*., 1998; Huang *et al*., 1999; Fagiolini & Hensch, 2000).

During early development, several studies have characterized the spatiotemporal dynamics of KCC2 and NKCC1 expression across brain regions in rodents and humans. In the rodent brain, KCC2 expression emerges postnatally in a region-specific pattern, with early expression in brainstem, spinal cord, and paleocortical regions, followed by delayed upregulation in neocortical and hippocampal areas (Stein *et al*., 2004; Takayama & Inoue, 2010; Kovács *et al*., 2014; Zavalin *et al*., 2024). Subcellular distribution studies further indicate layer-specific and isoform-specific regulation, with KCC2b predominating in cortical regions during postnatal development (Uvarov *et al*., 2009; Markkanen *et al*., 2014). In parallel, NKCC1 expression peaks during early postnatal stages and progressively declines during this period (Marty *et al*., 2002; Ludwig *et al*., 2003; Dzhala *et al*., 2005). In the human brain, KCC2 expression increases progressively throughout late gestation and continues to rise postnatally into the first year of life, suggesting conserved developmental dynamics across species (Sedmak *et al*., 2016).

Recent evidence shows that early-life caregiving experiences can modulate the developmental trajectory of these cotransporter expressions. Specifically, maternal separation increases NKCC1 expression and delays KCC2 upregulation in the hippocampus (Furukawa *et al*., 2017; Hu *et al*., 2017). Conversely, pharmacological blockade of NKCC1 can prevent these changes (Hu *et al*., 2017). Additionally, prolonged maternal separation accelerated the switch of GABA without changes in KCC2 expression, but by reducing NKCC1 cotransporter sensitivity to bumetanide in a sex-specific manner in the hippocampus (Galanopoulou, 2008). Conversely, exposure to the limited bedding and nesting (LBN) paradigm accelerates the GABA switch in the prefrontal cortex by reducing NKCC1 levels without altering KCC2 (Karst *et al*., 2023). These findings suggest that early maternal caregiving influences the maturation of inhibitory, with chloride transporter expression acting as a structural correlate of experience-dependent regulation. However, it remains unclear whether such modulation also occurs in brain regions that are critical for infant attachment learning, such as the piriform cortex.

The piriform cortex plays a central role in infant attachment learning in rodents. This region supports early odor preference learning through synaptic plasticity that occurs within a sensitive postnatal window (Sullivan, 2003; Yuan *et al*., 2014; Colombel *et al*., 2023). During natural rearing, pups acquire this learning during mother-infant interactions in the nest, expressed as a strong preference for the motheŕs odor (Leon, 1992; Sullivan, 2003). When this interaction is disrupted, this preference can be diminished (Perry *et al*., 2019). Given its role in attachment learning and its heightened synaptic plasticity during the early postnatal stage, the piriform cortex represents an ideal target for investigating how maternal caregiving experiences might influence the maturation of inhibitory circuits and their potential molecular substrate, such as the chloride cotransporters KCC2 and NKCC1.

Our previous work has shown that during the early postnatal period, from the sensitive period of attachment learning, pups younger than postnatal (P) 10 to the postsensitive period (> P10), GABAergic inputs to the anterior piriform cortex (aPC) pyramidal neurons mature from less frequent to faster synaptic activity, and the GABA reversal potential shifts from more depolarized to more hyperpolarized values (Pardo, 2018; Pardo *et al*., 2018). We also found that layer 2/3 pyramidal neurons exhibit intrinsic maturation, transitioning from a more depolarized resting membrane potential to a hyperpolarized state, along with changes in input-dependent activity (Oruro *et al*., 2020a). By integrating these data into a computational model of the olfactory bulb- piriform cortex circuit, we demonstrated that GABAergic signaling remains depolarizing in developing aPC pyramidal neurons, enhancing the excitatory drive and facilitating rapid odor learning during the sensitive period for attachment (Oruro *et al*., 2020b). Because the polarity of GABA signaling is determined by the chloride gradient, which is regulated by the developmental state of KCC2 and NKCC1, the depolarizing action we observed in aPC neurons is likely dependent on the maturation of these cotransporters. This early excitatory action of GABA, which in other brain regions is supported by the developmental state of KCC2 and NKCC1, suggests that similar maturational changes may occur in the piriform cortex during the sensitive period of attachment and could be sensitive to variations in maternal caregiving experience.

Although KCC2 expression in the three-layered piriform cortex is detectable from birth, initially confined to dendrites in the superficial layers at P0-P3, extending to deep layers by P4, and by P6 showing increased intensity across all layers with a pattern that remains stable through P14 (Kovács *et al*., 2014), the precise parallel developmental trajectories of KCC2 and NKCC1 during early stages and whether they are modulated by maternal caregiving remains unknown in this region, particularly during the sensitive period for attachment learning. Here, we address this gap by first characterizing the expression profiles of KCC2 and NKCC1 in the piriform cortex across the first three postnatal weeks, comparing male and female expression to adult levels. Second, we evaluate whether LBN exposure from P2-P9 alters transporter expression at P15 in both anterior (aPC) and posterior piriform cortex (pPC). The maternal behavior of the animals used in this study was previously characterized, revealing marked alterations in the structure and temporal organization of care (Pardo *et al*., 2024). Although LBN P15 pups show attachment behavior comparable to controls in home-nest preference and nipple-attachment tests, they exhibited more isolation-induced ultrasonic vocalizations below 60 kHz (Pardo *et al*., 2025). Finally, to explore how the observed expression profiles of chloride cotransporters shape inhibitory circuit function and influence excitability in developing aPC pyramidal neurons, we implemented a Hodgkin-Huxley computational model with dynamic ion concentrations (Lemaire *et al*., 2021) adapted for developing aPC pyramidal neurons.

## 2. Materials and Methods

### 2.1. Experiment 1

#### 2.1.1. Animals

Pregnant Sprague Dawley rats (Charles River Laboratories) were maintained in the animal facility at the Universidad Andina del Cusco (located at 3300 meters above sea level). On gestational days 17-18, pregnant rats were individually housed in standard cages (30 x 30 x 18 cm) with abundant bedding material for nest construction. The day of parturition was designated as postnatal day (P) 0. On P2, litters were culled to eight pups each, with a balanced sex ratio when possible. Pups remained with their mothers in their home cages under a 12h light/dark cycle (lights on at 6:00 A.M.), with controlled temperature (22-23°C) and humidity (45%). Food and water were available *ad libitum*. Litters were left undisturbed, with minimal handling every five days for cage cleaning and replenishment of food and water. All procedures were conducted in accordance with the NIH Guide for Animal Care and Use of Laboratory Animals and were approved by the Institutional Ethics Committee (CIEI) of the Andean University of Cusco (N°002- 2022-CIEI-UAC).

#### 2.1.2. Tissue collection and preparation

Whole piriform cortex tissue was collected at P5-6, P7-8, P11-12, P13-14, P15-16, P19- 20, P21-22, and adults. For each age, two pups per litter were euthanized and dissected on ice using the rhinal fissure and optic chiasm as landmarks. Tissue was stored at - 80°C until analysis (see detailed description in **Supplementary Material**, **subsection 1.1**).

#### 2.1.3. Western blot

Protein expression levels of KCC2 and NKCC1 in the piriform cortex were assessed by Western blot. Tissue samples were lysed in mammalian lysis buffer supplemented with protease inhibitors, and total protein concentrations were quantified using the Bradford method. Equal amounts of protein (200 µg per sample) were separated on SDS-Polyacrylamide gels, transferred to PVDF membranes, and incubated with primary antibodies against KCC2, NKCC1, and β-actin as a loading control. Immunoreactive bands were visualized using DAB substrate and quantified using ImageJ software. Details of buffer compositions, antibody specifications, and detection procedures are provided in the **Supplementary Material (subsections 1.2-1-3).**

All protein expression levels were normalized to β-actin and expressed as percentages relative to the mean values of adult rats, enabling developmental comparison across postnatal time points (P5–8, P11–14, P15–18, and P19–22).

#### 2.1.4. Quantitative RT-PCR

Total mRNA from piriform cortex samples was extracted with TRIzol and reverse-transcribed into cDNA. Quantitative PCR was performed using SYBR Green detection chemistry (Applied Biosystems, Canada) on an AriaMX Real-Time PCR System (Agilent, USA). Expression of KCC2 (Slc12a5), NKCC1 (Slc12a2), and housekeeping genes GAPDH and Hprt1 was quantified using gene-specific primers (see **Table S1, Supplementary Material**). Relative expression levels were calculated using the ΔΔCq method, normalized to the geometric mean of GAPDH and Hprt1. Detailed reaction conditions, primer efficiencies, and validation procedures are reported in the **Supplementary Material (subsection 1.4)**.

All mRNA expression levels were expressed as percentages relative to the mean adult rat (P134) values, enabling developmental comparison across postnatal time points (P5– 8, P11–14, P15–18, and P19–22).

#### 2.1.5. Immunofluorescence

To examine the cellular localization of KCC2 and NKCC1 in the piriform cortex across postnatal development, immunofluorescence staining was performed on coronal brain sections containing the anterior (aPC) and posterior (pPC) piriform cortex obtained at P5, P14, P22, and adult. The brains of three naïve animals per sex were collected. The full protocol is described in the **Supplementary Material (subsection 1.5).** This procedure aimed to qualitatively verify the presence of KCC2 and NKCC1 in the neurons of the piriform cortex, aPC, and pPC. Fluorescence intensity was not quantified; instead, the expression of each cotransporter was assessed based on its co- localization with NeuN-positive cells. Merged fluorescence signals indicating overlap between transporter and neuronal markers were used to confirm neuronal expression.

This approach provided cell-type–specific spatial context for interpreting the whole-tissue expression data obtained from Western blot and qRT-PCR.

#### 2.1.6. Statistical analysis

The effect of postnatal age on gene and protein expression (RT-qPCR and Western blot assays) was analyzed using one-way ANOVA or Kruskal-Wallis tests, as appropriate, followed by post hoc comparison. Data are presented as mean ± SEM or median (IQR), depending on the datás normality. Sample sizes (N) are detailed in figure legends. All statistical analysis and data visualization were performed with GraphPad Prism version 9.1.2 (San Diego, CA, USA), with *p*<0.05 considered statistically significant. Asterisks indicate significant group differences in tables and figures.

### 2.2. Experiment 2

#### 2.2.1. Animals and early-life stress paradigm

A total of 12 litters of Sprague Dawley rats were used in this experiment. Animals were maintained under the same housing conditions as described in Experiment 1 and were handled in accordance with the guidelines and regulations approved by the Institutional Ethics Committee (CIEI), as referenced in Experiment 1. From PND 2 to 9, litters were exposed to the limited bedding and nesting (LBN) protocol, as described (Pardo *et al*., 2024). Pups included in the present study were the offspring of the dams assessed in that previous study. Briefly, on PND 2, litters were culled to eight pups each (four males and four females when possible). Litters assigned to the LBN condition were transferred to cages with limited bedding material and a wire mesh floor, where they remained until the morning of PND 10. Litters assigned to the control condition were placed in standard cages with abundant nesting and bedding material. After the end of the LBN protocol, control and LBN litters were housed under standard conditions with abundant bedding. Litters were left undisturbed until PND 15, when tissue collection was performed.

#### 2.2.2. Tissue collection and preparation

On PND 15, one male and one female from each litter were randomly selected and removed from their home cages. Pups were euthanized following the same method described in Experiment 1. Brains were extracted following the protocol described in our previous study (Medina-Saldivar *et al*., 2024), immediately wrapped in labeled aluminum foil, snap-frozen on dry ice, and stored at -80°C. Brains were coronally sectioned at 500 μm for anatomical dissection, using a cryostat (CM 1520, Leica, Germany) maintained at -10°C. Sections were mounted on gelatin-coated slices placed on a -20°C platform within the cryostat. Tissue micropunches of the aPC and pPC were obtained using a 1.0 mm microdissector (EMS Aluminum Punch Kit, Electron Microscopy Science, USA), guided by anatomical coordinates from the developing rat brain atlas (Khazipov *et al*., 2015) (aPC: approx. Bregma 1.00 mm to -0.5 mm; pPC: approx. Bregma -1.20 mm to -3.20 mm) for infant rats and for adult rats approximately equal to Paxinos rat brain atlas (aPC: aprox Bregma 1.70 mm to 0.2 mm; pPC: aprox Bregma -1.88 mm to -3.88 mm) (Paxinos & Charles Watson, 2007). Micropunched samples from both hemispheres were pooled for each region and stored in RNase-free 1.5 mL microtubes at -80°C until RNA extraction were performed.

#### 2.2.3. RNA extraction

Total RNA was extracted from pooled bilateral micropunches of aPC and pPC. The protocol followed was the same as in Experiment 1, with the addition of Ribolock RNase Inhibitor (Thermo Scientific, E00382) during the extraction step to preserve RNA integrity. Samples were stored at -80°C until reverse transcription and further analysis.

#### 2.2.4. Western Blot and Quantitative RT-PCR

Protein and mRNA expression levels of KCC2 and NKCC1 were analyzed separately for aPC and pPC using the same protocols described in Experiment 1 for Western blot and RT-qPCR. No quantification of fluorescence intensity was performed for these samples; instead, comparison focused on expression levels across conditions and regions.

For both protein and gene expression analysis, values were first normalized to the internal reference genes (β-actin for Western blot and the geometric mean of Gapdh and Hprt1 for qPCR). These normalized values were then expressed as percentages relative to the mean of the control group, set as 100%, enabling direct comparison of LBN-exposed animals to baseline control levels.

#### 2.2.5. Statistical analysis

The effects of early-life stress on protein and gene expression levels (assessed by Western Blot and RT-qPCR assays) were analyzed using a one-sample *t* test. Data are reported as mean ± SEM . The number of subjects per group (N) is indicated in figure legends. All statistical analysis and data visualization were performed using GraphPad Prism version 9.1.2 (San Diego, CA, USA). Statistical significance was set at *p*<0.05 and significant group differences are indicated by asterisks in figures and tables.

### 2.3. Computational modeling and simulation

To explore the functional consequences of the observed KCC2 expression profiles, we implemented a single-cell Hodgkin-Huxley type model with dynamic ion concentrations, adapted from (Lemaire et al, 2021), to simulate the activity of a postsynaptic pyramidal neuron in the aPC during early postnatal development (**Figure S1, Supplementary Material**). Pyramidal neurons in the aPC are projection neurons that receive olfactory input from the olfactory bulb as well as recurrent excitatory projections from other pyramidal neurons and inputs from different brain regions (Oruro *et al*., 2020a)(**Figure S1A, Supplementary Material**). Ion dynamics were modeled through coupled intra-and extracellular concentrations, ion currents, and synaptic activation (**Figure S1B, Supplementary Material**).

The model included voltage-gated sodium (Na^+^), potassium (K^+^), and calcium (Ca^2+^) channels, the Na^+^/K^+^ ATPase pump, chloride cotransporters (NKCC1 and KCC2), and ionotropic receptor channels relevant for GABAergic synaptic transmission, all parametrized according to (Lemaire *et al*., 2021). In contrast to the original model, we excluded glutamatergic synapses and GABAergic interneurons, retaining only postsynaptic GABAA receptor conductance to simulate inhibitory input. Cotransporter dynamics were standardized to the resting membrane potential of aPC pyramidal neurons at P15 (-55.54 mV), based on whole-cell voltage-clamp data from our previous work (Oruro *et al*., 2020a).

The temporal evolution of the model is governed by a system of differential equations adapted from (Lemaire *et al*., 2021), with those specific to GABAergic neurons excluded. To simulate the effects of LBN, cotransporter activity was parametrized using our Western Blot results for KCC2 and NKCC1 expression.

The system of differential equations was implemented in NetLogo 6.6.2 software (Wilensky, 1999; http://ccl.northwestern.edu/netlogo) using explicit Euler integration (time step=0.01ms) to approximate solutions to the ordinary differential equations. All differential equations from (Lemaire *et al*., 2021) were adapted, with minor modifications regarding the inhibitory neuron contribution (details in **Table S2,**

**Supplementary Material).** All state variables and parameters are provided in the **Supplementary Material (section 2)**, and the simulation code and data are available in https://sites.google.com/view/orurolab/datapapercotransportadores.

## 3. Results

### 3.1. Ontogeny of expression patterns of KCC2 in male and female naïve rats

To obtain an overview of KCC2 expression in the developing piriform cortex, we analyzed KCC2 protein and mRNA levels in whole-tissue homogenates obtained from the entire piriform cortex of male and female rats at different postnatal stages, relative to adult animals (**Figure 1 and Figure 2**). Male and female protein and mRNA expression levels were normalized to those of adults of the same sex; therefore, we did not provide a sex comparison in this study.

**Figure 1.**
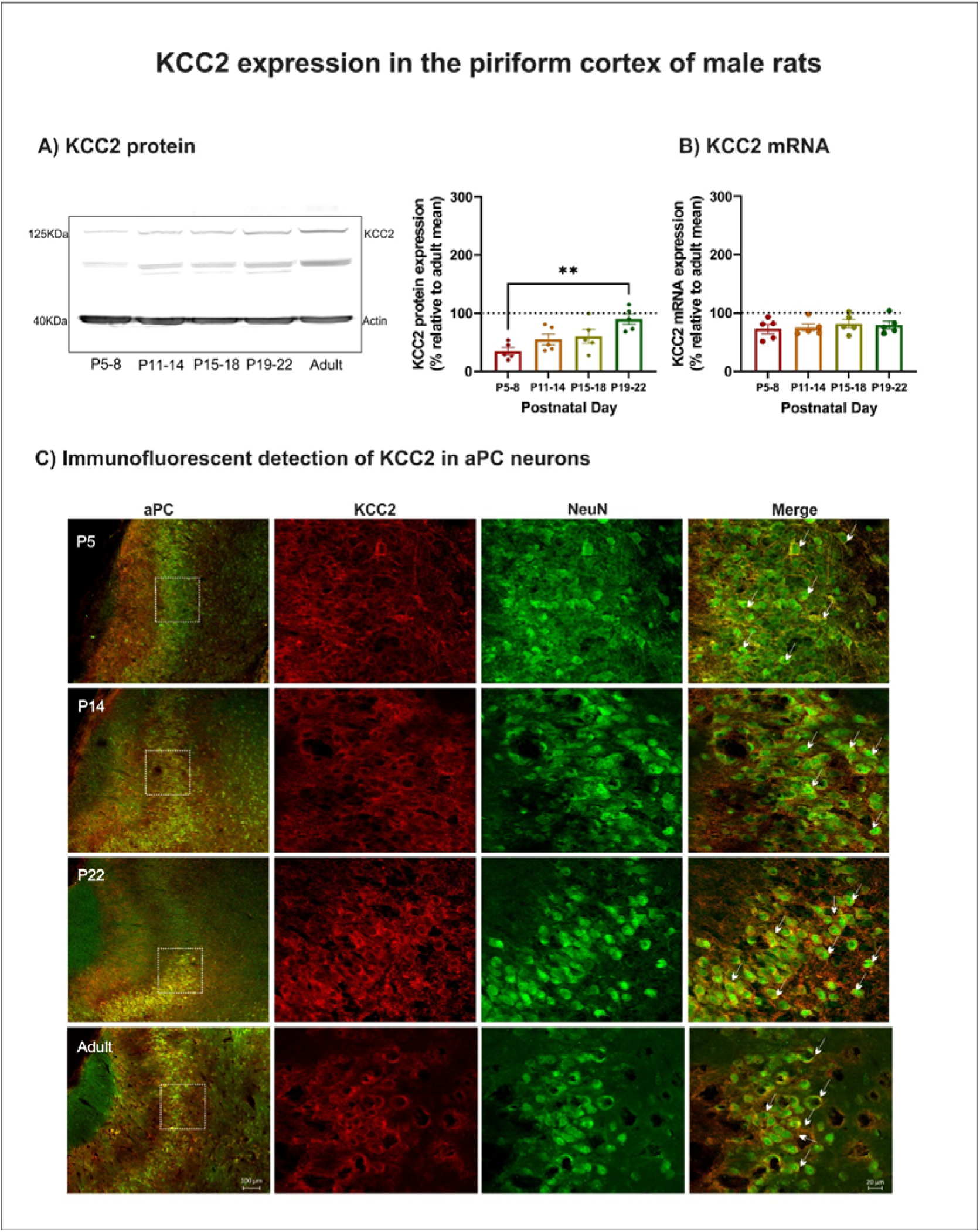
KCC2 expression patterns in the piriform cortex of male rats during early postnatal development. **A**) Representative Western blot showing KCC2 protein levels in the piriform cortex across four developmental stages: P5-8, P11-14, P19-22, and adult. The bar graph depicts relative KCC2 expression as a percentage of the adult value, indicated by the dashed line at 100%. **B)** Relative KCC2 mRNA levels (fold change % of adult) at P5-8, P11-14, P19-22. **C**) Immunofluorescence images of the anterior piriform cortex (aPC) at P5, P14, P22, and adult. Overview images (left column) show the aPC region (scale bar=100 µm); magnified views show KCC2 (red), NeuN (green), and merged signals in layer 2/3 (scale bar= 20 µm). White arrows in merged panels indicate neurons co-expressing KCC2 and NeuN. Data in panels A and B are presented as mean ± SEM. Statistical comparisons were performed using one-way ANOVA followed by post hoc Tukey’s multiple comparisons test (n=4-5 per group). \**p*<0.05, \*\**p*<0.01, \*\*\**p*<0.001, \*\*\*\**p*<0.0001.

**Figure 2.**
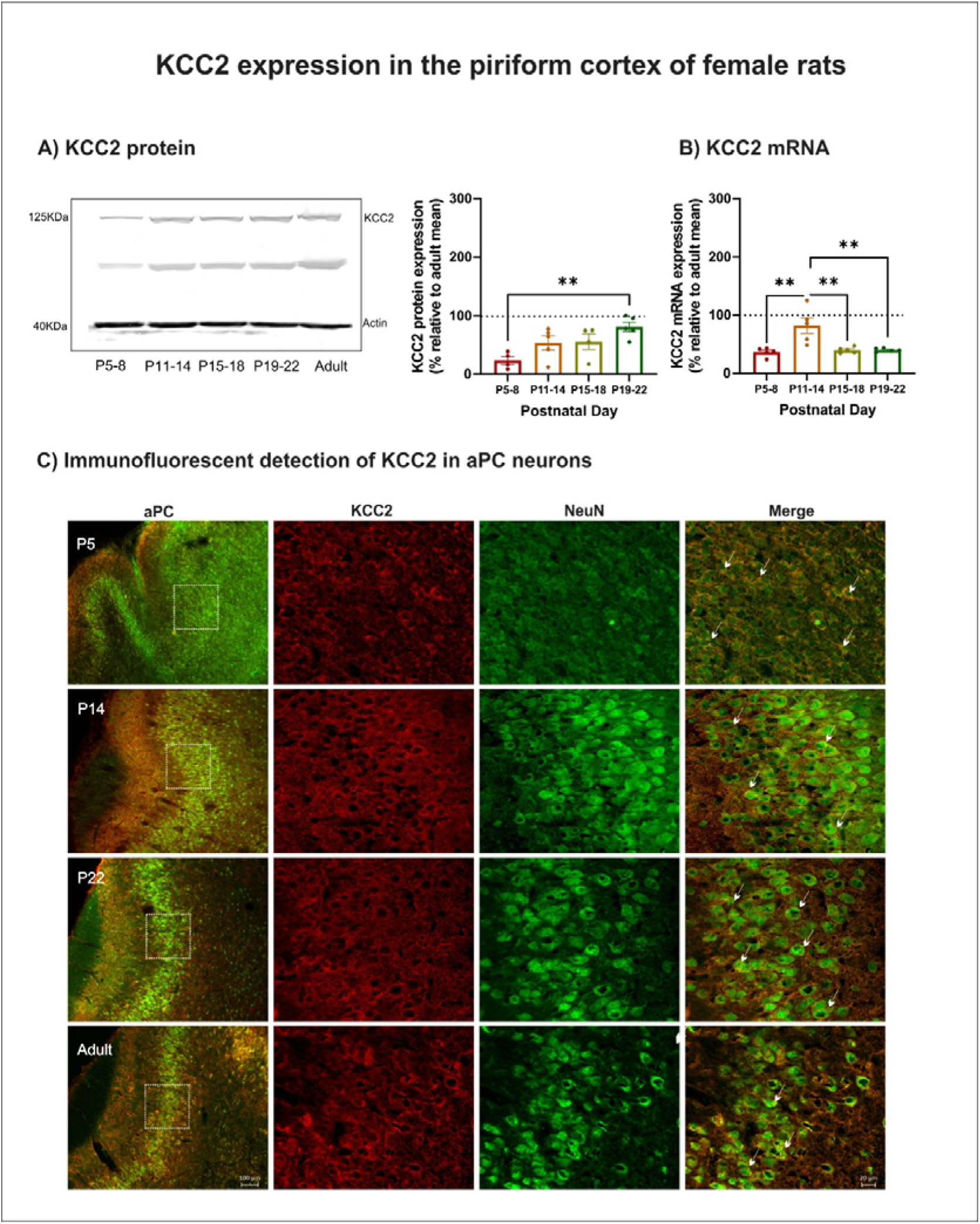
KCC2 expression patterns in the piriform cortex of female rats during early postnatal development. **A**) Representative Western blot showing KCC2 protein levels in the piriform cortex across four developmental stages: P5-8, P11-14, P19-22, and adult. The bar graph depicts relative KCC2 expression as a percentage of the adult value, indicated by the dashed line at 100%. **B)** Relative KCC2 mRNA expression (fold change % of adult) at P5-8, P11-14, P19-22. **C**) Immunofluorescence images of the anterior piriform cortex (aPC) at P5, P14, P22, and adult. Overview images (left column) show the aPC region (scale bar=100 µm); magnified views show KCC2 (red), NeuN (green), and merged signals in layer 2/3 (scale bar= 20 µm). White arrows in merged panels indicate neurons co-expressing KCC2 and NeuN. Data in panels A and B are presented as mean ± SEM. Statistical comparisons were performed using one-way ANOVA followed by post hoc Tukey’s multiple comparisons test (n=4-5 per group). \**p*<0.05, \*\**p*<0.01, \*\*\**p*<0.001, \*\*\*\**p*<0.0001.

***Males***. As shown in **Figure 1A**, KCC2 protein levels in male rats increased progressively with age. A one-way ANOVA revealed a significant effect of age on KCC2 protein levels [F_(3,16)_ = 5.848, *p* = 0.0068], with post hoc Tukey analysis indicating that expression at P19–22 was significantly higher than at P5–8 (*p* = 0.0038). In contrast, analysis of KCC2 mRNA expression (fold change relative to adult levels) showed no significant effect of age [F_(3,16)_=0.3236, *p*=0.8082] (**Figure 1B**).

Immunofluorescence staining (**Figure 1C**) confirmed that KCC2 was primarily expressed in NeuN-positive neurons of the aPC, with strong localization in layer 2/3, as indicated by co-localization with NeuN. A qualitative increase in KCC2 labeling intensity was observed with age, consistent with protein expression data. These immunofluorescence staining were performed in a separate cohort of pups, using coronal brain slices to allow regional and laminar resolution of KCC2 expression.

***Females***. A similar developmental pattern was observed in female rats (**Figure 2**). KCC2 protein levels increased significantly with age [F_(3,14)_ = 4.880, *p* = 0.0158], with higher expression at P19–22 compared to P5–8 (**Figure 2A**). In contrast to males, however, KCC2 mRNA expression in females also showed a significant effect of age [F_(3,16)_ = 9.337, *p* = 0.0008]. Post hoc comparisons revealed that mRNA expression at P11–14 was significantly higher than at P5–8 (*p* = 0.0017), and that expression at P15– 18 and P19–22 was significantly higher than at P11–14 (*p* = 0.0034 and *p* = 0.0035, respectively) (**Figure 2B**). Immunofluorescence results confirmed strong KCC2 labeling in layer 2/3 aPC neurons (**Figure 2C**), with a clear increase in signal intensity across developmental stages.

KCC2 expression was also observed in pPC neurons in both male (**Figure S2A**) and female rats (**Figure S3A**). Together, these results indicate a progressive postnatal increase in KCC2 protein in both sexes, also suggesting a sex-dependent difference in mRNA regulation.

### 3.2. Ontogeny of expression patterns of NKCC1 in male and female naïve rats

Developmental trajectory of NKCC1 expression in the piriform cortex was also examined at protein and gene levels in whole-tissue homogenates from the entire piriform cortex of male and female rats from P5–8 to P19–22 (**Figures 3 and 4**).

**Figure 3.**
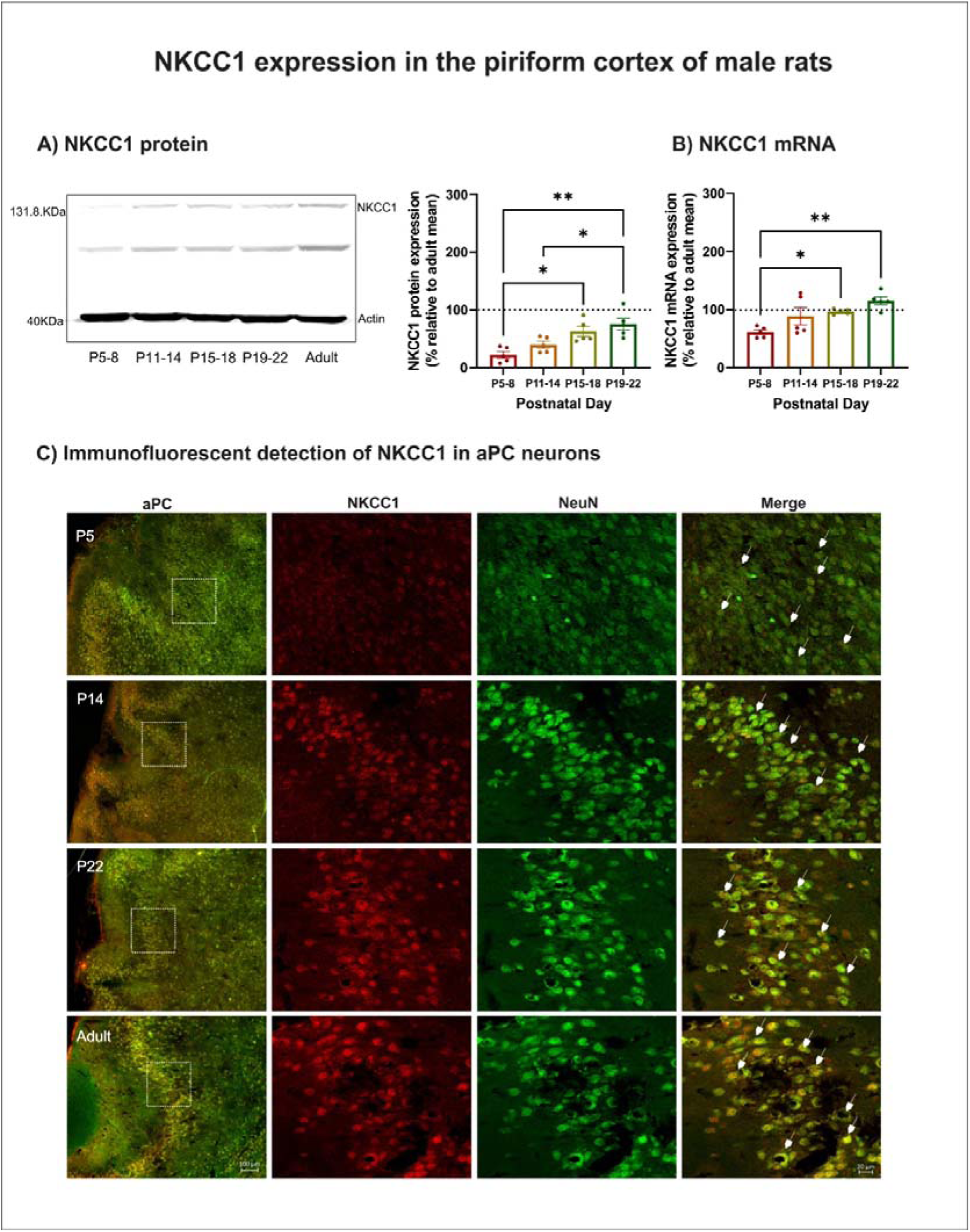
NKCC1 expression patterns in the piriform cortex of male rats during early postnatal development. **A**) Representative Western blot showing NKCC1 protein levels in the piriform cortex across four developmental stages: P5-8, P11-14, P19-22, and adult. The bar graph depicts relative NKCC1 expression as a percentage of the adult value, indicated by the dashed line at 100%. **B)** Relative NKCC1 mRNA expression (fold change % of adult) at P5-8, P11-14, P19-22. **C**) Immunofluorescence images of the anterior piriform cortex (aPC) at P5, P14, P22, and adult. Overview images (left column) show the aPC region (scale bar=100 µm); magnified views show NKCC1 (red), NeuN (green), and merged signals in layer 2/3 (scale bar= 20 µm). White arrows in merged panels indicate neurons co-expressing NKCC1 and NeuN. Data in panels A and B are presented as mean ± SEM. Statistical comparisons were performed using one-way ANOVA followed by post hoc Tukey’s multiple comparisons test (n=4-5 per group). \**p*<0.05, \*\**p*<0.01, \*\*\**p*<0.001, \*\*\*\**p*<0.0001.

**Figure 4.**
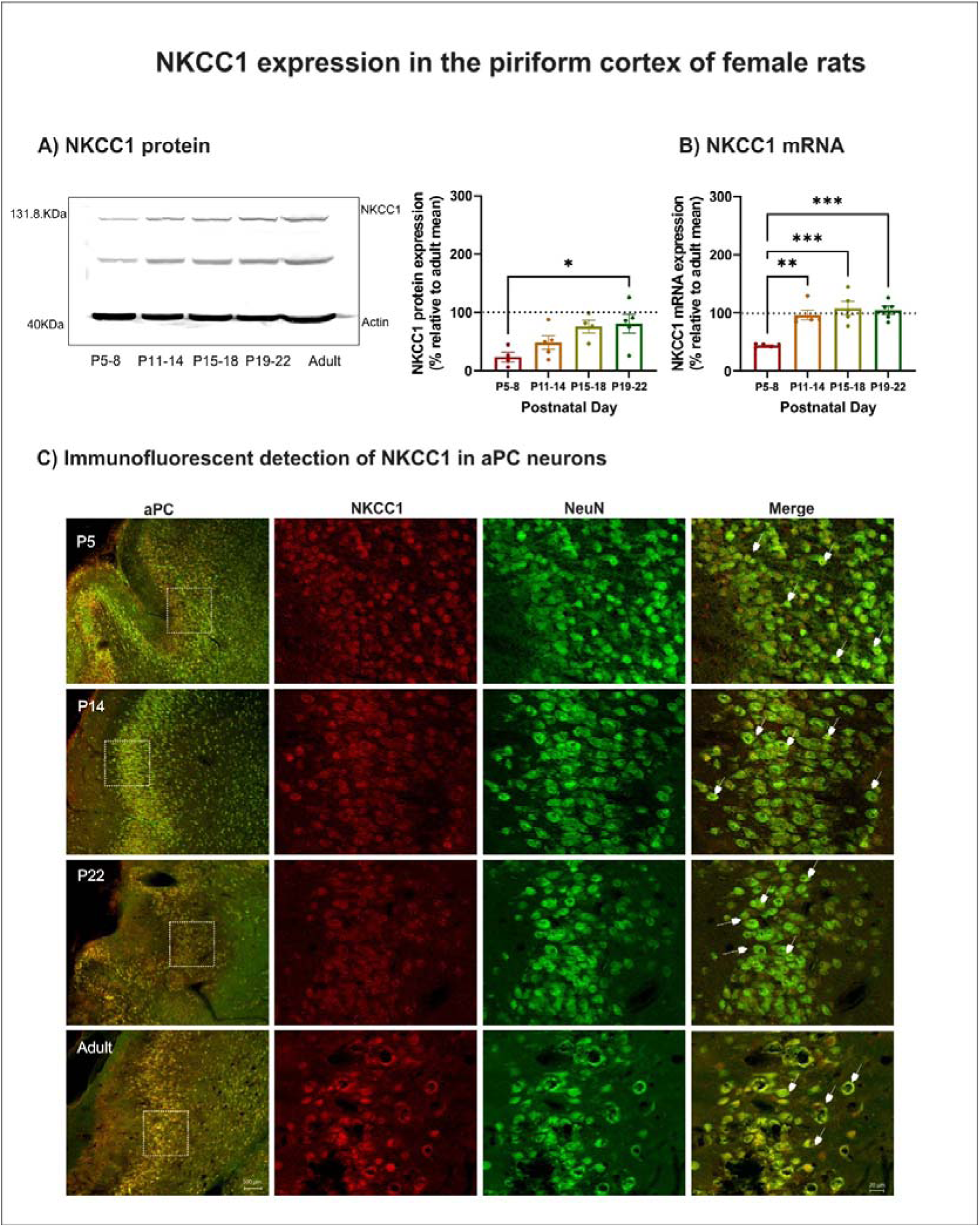
NKCC1 expression patterns in the piriform cortex of female rats during early postnatal development. **A**) Representative Western blot showing NKCC1 protein levels in the piriform cortex across four developmental stages: P5-8, P11-14, P19-22, and adult. The bar graph depicts relative NKCC1 expression as a percentage of the adult value, indicated by the dashed line at 100%. **B)** Relative NKCC1 mRNA expression (fold change % of adult) at P5-8, P11-14, P19-22. **C**) Immunofluorescence images of the anterior piriform cortex (aPC) at P5, P14, P22, and adult. Overview images (left column) show the aPC region (scale bar=100 µm); magnified views show NKCC1 (red), NeuN (green), and merged signals in layer 2/3 (scale bar= 20 µm). White arrows in merged panels indicate neurons co-expressing NKCC1 and NeuN. Data in panels A and B are presented as mean ± SEM. Statistical comparisons were performed using one-way ANOVA followed by post hoc Tukey’s multiple comparisons test (n=4-5 per group). \**p*<0.05, \*\**p*<0.01, \*\*\**p*<0.001, \*\*\*\**p*<0.0001.

***Males***. NKCC1 protein levels in male rats showed a significant age-related increase [F_(3,16)_=9.049, *p*=0.0010], with post hoc comparisons indicating significantly higher expression at P15–18 and P19–22 compared to P5–8 (*p* = 0.0011 and *p* = 0.0108, respectively), and at P19–22 compared to P11–14 (*p* = 0.0257) (**Figure 3A**). Paralleling protein, NKCC1 mRNA levels also increased significantly with age [F_(3,16)_=6.987, *p*= 0.0032], with levels at P15–18 and P19–22 higher than at P5–8 (*p* = 0.0019 and *p* = 0.0420, respectively) (**Figure 3B**). Immunofluorescence staining (**Figure 3C**) showed NKCC1 in aPC neurons, with predominant localization in layer 2/3 and a visible increase in labeling intensity across developmental stages.

***Females***. NKCC1 protein levels in female rats also increased significantly with age [F_(3,14)_ = 4.175, *p* = 0.0263], with higher levels at P19–22 compared to P5–8 (*p* = 0.0308) (**Figure 4A).** mRNA levels also increased as a function of age [F_(3,16)_ = 13.81, *p* = 0.0001], with significantly higher NKCC1 mRNA levels at P11–14, P15–18, and P19–22 compared to P5–8 (*p*=0.0015, *p*=0.0002, and *p*=0.0004, respectively) (**Figure 4B**). Immunofluorescence staining (**Figure 4C**) followed this tendency, with NKCC1 expression in NeuN-positive neurons increasing across postnatal stages.

NKCC1 was also detected in pPC neurons of male (**Figure S2B**) and female rats (**Figure S3B**). Together, these results indicate a progressive upregulation of NKCC1 protein and mRNA levels across postnatal development, with an earlier onset in females.

### 3.3. Influence of early maternal caregiving alterations on KCC2 and NKCC1 levels

To assess the impact of early caregiving alterations on KCC2 and NKCC1 levels, we analyzed protein and mRNA levels in micropunch samples obtained from the aPC and pPC of P15 male rats exposed to the LBN paradigm during the first postnatal week (**Figure 5**). This age was chosen because it coincides with a stage when pups begin to exhibit increased behavioral independence (Thiels *et al*., 1990) and are capable of discriminating their motheŕs odor from cues (Kojima & Alberts, 2009), making it particularly relevant for evaluating the potential neurodevelopmental consequences of altered early caregiving.

**Figure 5.**
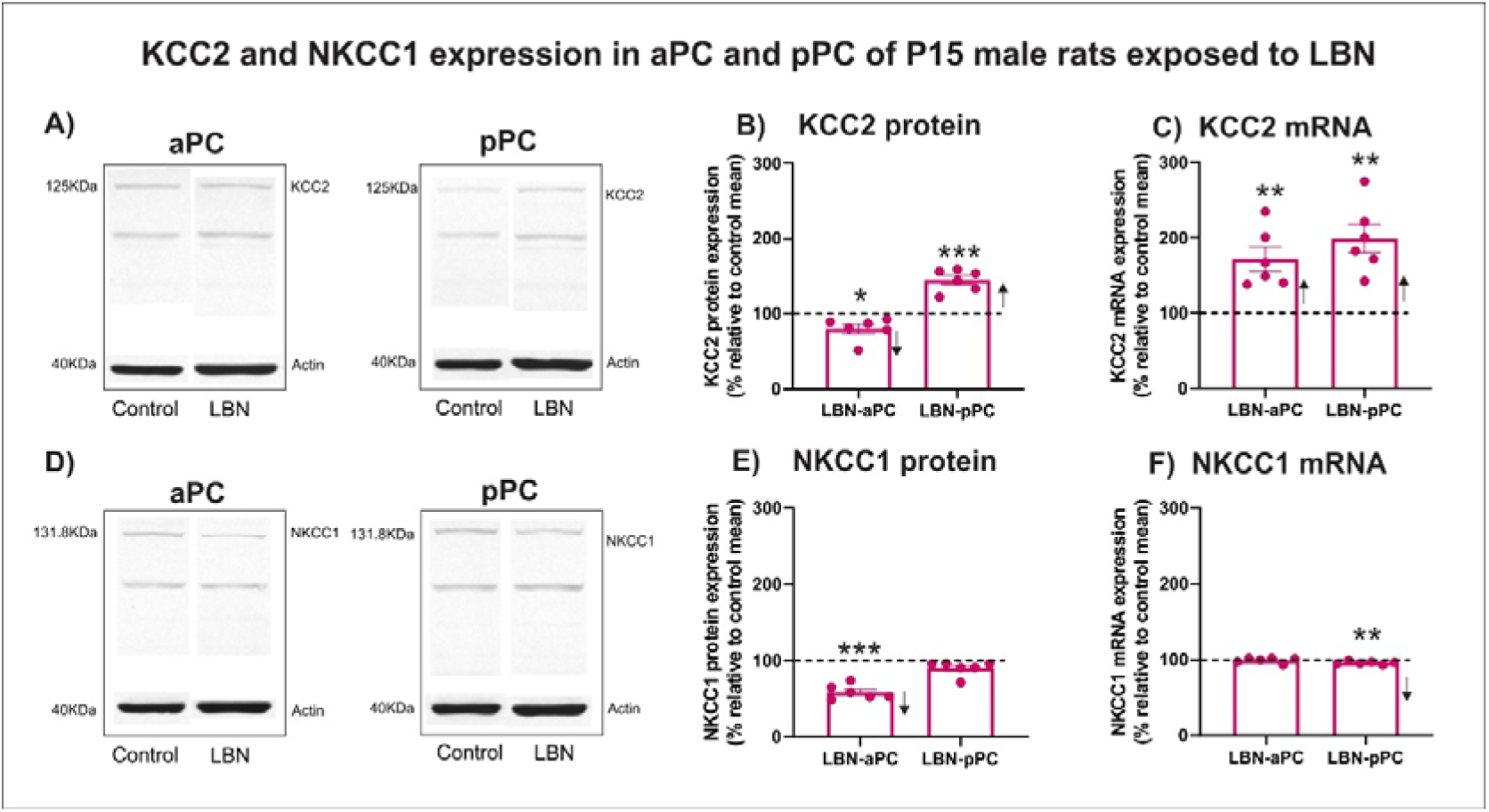
KCC2 and NKCC1 expression in the aPC and pPC of P15 male rats exposed to LBN. **A**) Representative Western blots showing KCC2 protein expression in aPC and pPC of control and LBN pups. **B**) Quantification of KCC2 protein levels in aPC and pPC of LBN-exposed pups expressed as a percentage relative to the control group mean. **C**) KCC2 mRNA expression in aPC and pPC of LBN-exposed pups, expressed as percentage relative to the control group mean. **D**) Representative Western blots showing NKCC1 protein expression in aPC and pPC of control and LBN pups. **E**) Quantification of NKCC1 protein levels in aPC and pPC of LBN-exposed pups expressed as a percentage relative to the control group mean. **F**) NKCC1 mRNA expression in aPC and pPC of LBN-exposed pups, expressed as a percentage relative to the control group mean. Data are presented as mean ± SEM (n=5-6 per group). Statistical comparisons were performed using one-sample t-test against a theoretical value of 100% (control group mean). \**p*<0.05, \*\**p*<0.01, \*\*\**p*<0.001, \*\*\*\**p*<0.0001.

The maternal behavior of the dams of these pups had been previously characterized, showing a marked increase in transitions between maternal repertories, particularly during the dark phase of the LBN exposure period (Pardo *et al*., 2024). These dynamic patterns, which reflect fragmentation and instability of maternal care, are summarized in **Figure S4**, which illustrates behavioral transitions networks and heatmaps across postpartum days 2, 5, and 9. Importantly, the pups used in this study showed no alterations in maternal odor preference (P13) or nipple attachment (P14), respectively (Pardo *et al*., 2025), suggesting that early attachment behavior, which relies on the ability of pups to learn the motheŕs odor (Sullivan, 2003), was not compromised. However, as shown in the following section, these early caregiving alterations were sufficient to induce changes at the molecular level in the expression of chloride cotransporters within the region that is necessary and sufficient for odor preference learning (Sullivan, 2003; Yuan *et al*., 2014; Colombel *et al*., 2023) .

***Males***. Male pups exposed to LBN showed significantly lower levels of KCC2 protein levels in the aPC, with values reaching 80.03±6.11% relative to the control value set at 100% [*t*(5)=3.267, *p*=0.0223, one-sample t-test analysis]. In contrast, in the pPC, KCC2 protein levels were significantly elevated [44.7[±[5.96%; *t*(5) = 7.508, *p* = 0.0007] (**Figure 5A, B**). KCC2 mRNA expression levels were also significantly increased relative to 100% in both aPC [171.5±15.86; *t*(6)=4.509, *p*=0.0063] and pPC [198.8±18.66; *t*(6)=5.295, *p*=0.0032] (**Figure 5,C**).

Regarding the NKCC1 cotransporter, protein levels in LBN male pups were significantly reduced in the aPC compared to 100% control levels [58.65±3.82%; *t*(5)=10.84, *p*=0.0001,one-sample t-test analysis] (**Figure 5 D,E**), but showed a slights reduction in the pPC [90.09±3.906%; *t*(59=2.538, *p*=0.0520] (**Figure 5 E**). NKCC1 mRNA levels were not significantly different from controls in aPC [ 99.41 ±1.405%; *t*(5)=0.4207, *p*=0.6915], but they were significantly reduced in the pPC [96.26±0.883%; *t*(5)=4.234, *p*=0.0082] ( **Figure 5,F**).

***Females***. Female pups exposed to LBN showed significantly lower levels of KCC2 protein levels in aPC, with values reaching 58.72±3.238% relative to the control value set at 100% [*t*(5)=12.75, *p<*0.0001]. Similarly, in the pPC, KCC2 protein levels were significantly reduced [33.88[±2.615%; *t*(5) = 25.29, *p<*0.0001] (**Figure 6A,B**). In contrast, KCC2 mRNA expression levels were significantly increased relative to 100% in both aPC [131.6±4.845; *t*(5)=6.516, *p*=0.0013] and pPC [135.1±2.595; *t*(5)=13.54, *p<*0.0001] (**Figure 6,C**).

**Figure 6.**
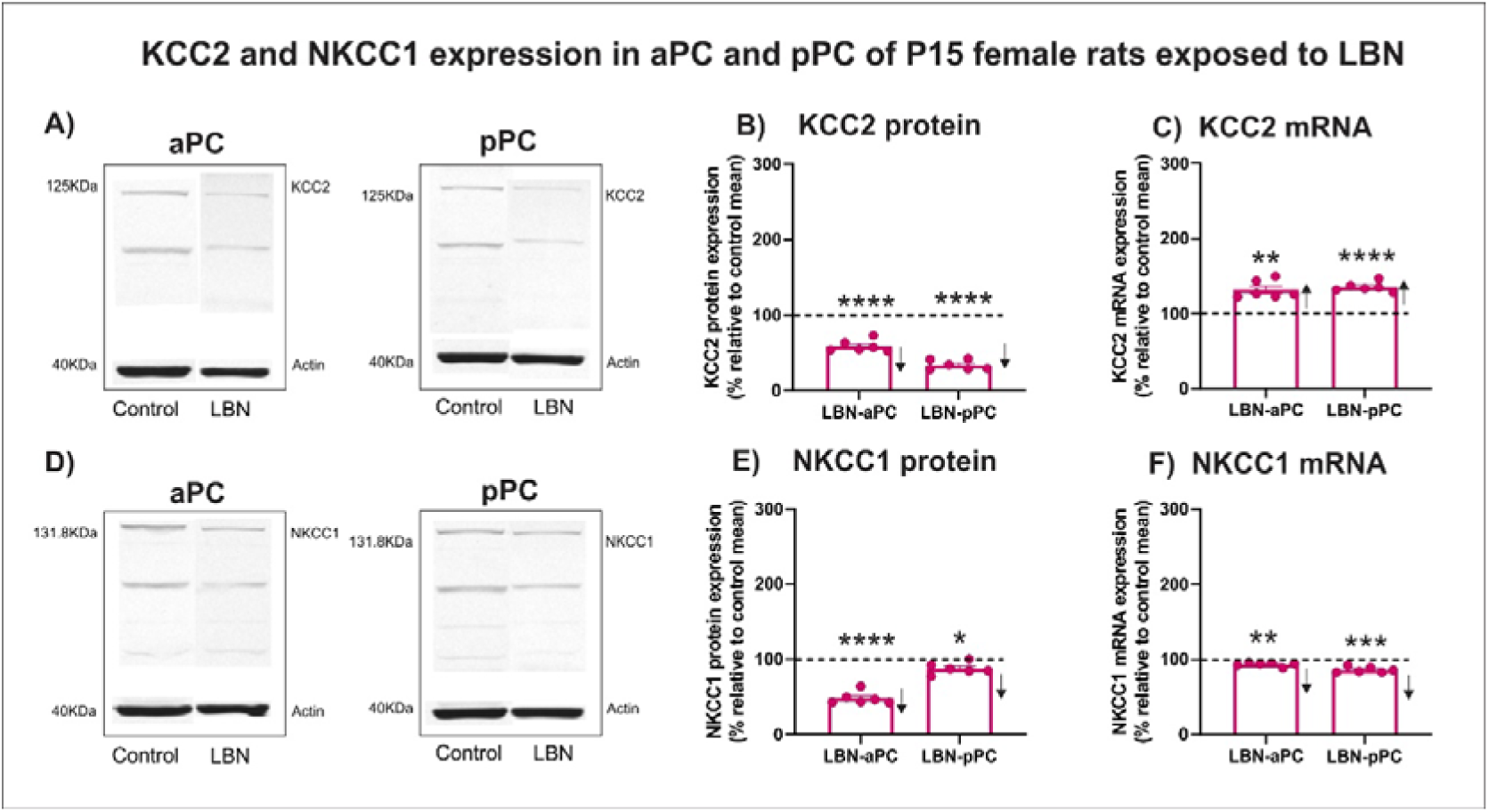
KCC2 and NKCC1 expression in the aPC and pPC of P15 female rats exposed to LBN. **A**) Representative Western blots showing KCC2 protein expression in aPC and pPC of control and LBN pups. **B**) Quantification of KCC2 protein levels in aPC and pPC of LBN-exposed pups expressed as a percentage relative to the control group mean. **C**) KCC2 mRNA expression in aPC and pPC of LBN-exposed pups, expressed as percentage relative to the control group mean. **D**) Representative Western blots showing NKCC1 protein expression in aPC and pPC of control and LBN pups. **E**) Quantification of NKCC1 protein levels in aPC and pPC of LBN-exposed pups expressed as a percentage relative to the control group mean. **F**) NKCC1 mRNA expression in aPC and pPC of LBN-exposed pups, expressed as a percentage relative to the control group mean. Data are presented as mean ± SEM (n=5-6 per group). Statistical comparisons were performed using one-sample t-test against a theoretical value of 100% (control group mean). \**p*<0.05, \*\**p*<0.01, \*\*\**p*<0.001, \*\*\*\**p*<0.0001.

Regarding the NKCC1 cotransporter, protein levels were significantly decreased in LBN female pups in both the aPC [48.27±3.39%; *t*(5)=15.26, *p<*0.0001] (**Figure 6 D,E**), and the pPC [87.66±3.169%; *t*(5)=3.894, *p*=0.0115] (**Figure 6 E**). NKCC1 mRNA levels were also significantly lower than control in both aPC [ 93.40±1.107%; *t*(5)=5.963, *p*=0.0019] and pPC [86.20±1.586%; *t*(5)=8.698, *p*=0.0003] ( **Figure 6 F**).

These results show that early caregiving alteration induced by the LBN paradigm impacts KCC2 and NKCC1 protein levels in the piriform cortex in a region-and sex-specific manner. In males, KCC2 protein levels increased in pPC, while in females it decreased in both regions. NKCC1 protein levels was consistently reduced in females but showed variable changes in males. At the transcriptional level, KCC2 mRNA increased in both sexes, while NKCC1 mRNA was reduced mainly in the pPC of LBN-exposed pups.

### 3.4. Computational simulation of KCC2 and NKCC1 expression impact on membrane potential

To explore how changes in the expression of chloride cotransporters at the molecular level could influence neuronal membrane dynamics, we implemented a computational model capable of linking these molecular changes to neuronal activity by integrating ion concentration dynamics and their effects on membrane potential. Based on a previous model (Lemaire et al, 2021), we implemented an HH model with dynamic ion concentration, allowing us to explore how the experimentally observed alterations in KCC2 and NKCC1 expression under LBN conditions could affect the membrane potential dynamics during GABAergic inputs.

#### 3.4.1. Dynamics of neuronal activity during GABAergic synaptic transmission

To evaluate how neuronal membrane activity changes during GABAergic synaptic transmission in the model, we first simulated the activity of a pyramidal neuron as a function of changes in KCC2 fraction, using a three-phase (A-B-A) protocol over a total period of 150000 ms (**Figure S5**). In this protocol, the same neuron underwent three sequential conditions across this time window. In the first condition (***before syn***), the neuron was simulated without activating the GABAergic synapse, from 0 ms to 70000 ms. In the second condition (***during syn***), the GABAergic synapse was activated every 25 ms, from 70001 to 140000 ms. In the final condition (***after syn***), the GABAergic synapse was deactivated, and the neuron was simulated from 140001 to 150000 to observe whether the model returned to resting conditions following synaptic transmission (**Figure S5, A**).

In all three conditions, the neuron was simulated across a range of KCC2 expression levels (expressed as fractions from 0.0 to 2.0). For each KCC2 fraction, simulations began with a baseline KCC2 value of 1.0 and a resting potential of -55.53 mV, reflecting the KCC2 expression level (100%) and the mean membrane potential observed in aPC pyramidal cells from our previous whole-cell current-clamp recordings in P15 rats (Oruro et al., 2020a), respectively. These values are represented as a red point in the graphs of **Figure S5**. Each simulation ran independently for every KCC2 fraction. Data were collected at the end of each time window: at 70000 ms for *before syn*, at 140000 ms for *during syn*, and at 150000 ms for *after syn*. The time window prior to GABAergic synapse activation was selected to ensure model stabilization to evaluate the potential excitatory effects of GABAergic input.

We analyzed resting membrane potential, ion reversal potentials, and intracellular ion concentrations of potassium, sodium, and chloride (**Figure S5**), as well as spiking activity and the relationship between membrane potential and chloride reversal potential (**Figure S5**).

***Before syn simulation***. Without GABAergic synapse activation, the membrane potential remained close to the resting membrane potential (-55.53 mV at KCC2 fraction 1.0). When the KCC2 fraction was reduced from 1.0 towards 0.0, membrane potential slightly depolarized, whereas increasing KCC2 above 1.0 led to a slight hyperpolarization (**Figure S5, B, *before syn*).** This was accompanied by a progressive decrease in potassium reversal potential, a slight increase in sodium reversal potential, and a marked increase in chloride reversal potential as the KCC2 fraction decreased toward 0.0; the opposite trends were observed as the KCC2 fraction increased toward 2.0 (**Figure S5, C, *before syn*)**. These changes occurred alongside an increase in intracellular potassium concentration, a slight decrease in intracellular sodium, and a more pronounced increase in intracellular chloride concentration at lower KCC2 fraction levels; again, the opposite pattern was observed as the KCC2 fraction increased beyond 1.0 (**Figure S5, D, *before syn*).** No spiking activity was observed at any KCC2 level (**Figure S6, A, *before syn***). These ionic and electrical dynamics were consistent with a slight reduction in the gap between the membrane potential and the chloride reversal potential curves (**Figure S6, B, *before syn*).**

***During syn simulation.*** Following GABAergic synapse activation, the membrane potential was hyperpolarized (approximately -58 mV), deviating from the resting potential (-55.53 mV at KCC2 fraction 1.0). As the KCC2 fraction decreased towards 0.0, this hyperpolarized state was maintained. In contrast, as the KCC2 fraction increased above 1.0, the membrane gradually depolarized, approaching resting levels and reaching approximately -56 mV when KCC2 was equal to 2.0 (**Figure S5, B, *during syn)***. The reversal potential for potassium, sodium, and chloride dropped markedly at KCC2 1.0. When the KCC2 fraction was reduced to below 1.0, the potassium reversal potential became more hyperpolarized, whereas increasing KCC2 above 1.0 led to less hyperpolarized values, even surpassing resting equilibrium values. Sodium reversal potential, on the other hand, increased towards its resting equilibrium, while chloride reversal potential became slightly more hyperpolarized, though it did not fully return to its resting values (**Figure S5, C, *during syn***). Intracellular potassium and chloride concentrations dropped sharply at KCC2 1.0 relative to resting levels, while sodium concentration increased. As KCC2 decreased toward 0.0, potassium remained low, sodium further increased, and chloride slightly increased. Conversely, as KCC2 increased beyond 1.0, potassium increased, sodium dropped dramatically, and chloride concentration remained relatively stable (**Figure S5, D, *during syn***). Neuronal activity increased during GABAergic stimulation, with 1403 spikes recorded at KCC2 1.0. This activity further increased as the KCC2 fraction decreased toward 0.0, reaching nearly 2000 spikes. In contrast, activity declined sharply as KCC2 increased beyond 1.0, with only a single spike at KCC2 fraction 2.0 (**Figure S6, A, *during syn*).** The increased activity at low KCC2 fractions coincided with a broader difference between the resting membrane potential and the chloride reversal potential (approximately 2.5 mV), while reduced activity at higher KCC2 fractions was supported by a narrowing gap, with both curves converging near -56 mV (**Figure S6, B, *during syn*).**

***After syn simulation.*** After GABAergic synaptic transmission was deactivated, resting membrane potential (**Figure S5, B, *after syn*)**, ion reversal potential (**Figure S5, C, *after syn*)**, and intracellular ion concentration (**Figure S5, D, *after syn*)** remained similar to those observed in the *during syn* condition. Similarly, spiking activity (**Figure S6, A, *after syn***) and the dynamics between membrane potential and chloride reversal potential (**Figure S6, B, *after syn***) closely resembled the previous phase. This suggests that the model does not immediately return to baseline after GABAergic input ceases, possibly indicating the need for a longer recovery period for ionic and electrical equilibrium.

Together, these results indicate that the model is capable of replicating key properties of early postnatal GABAergic transmission described in the literature. In particular, it captures the depolarizing effect of GABA when KCC2 expression is low, as typically observed during immature neurons, and the progressive reduction of this effect as KCC2 levels increase (Rivera *et al*., 1999). Beyond reproducing this classical shift in GABAergic polarity, the model also shows how membrane activity is shaped by coordinated changes in the reversal potential and intracellular concentrations of potassium, sodium, and chloride across different KCC2 fractions. These ionic dynamics provide a mechanistic substrate for understanding how KCC2 expression levels modulate excitability and recovery after GABAergic synaptic input, highlighting the value of this modeling approach to explore the non-linear and time-dependent interactions between ion homeostasis and synaptic transmission.

#### 3.4.2. LBN-induced changes in piriform cortex pyramidal neuron activity

Once the model was calibrated, we simulated hypotheses derived from our experimental findings on the effects of LBN on KCC2 and NKCC1 expression in aPC and pPC of P15 rat pups. To this end, we converted protein levels obtained from Western Blot analysis into fractions relative to expression levels observed in controls. Based on this, we simulated five biologically grounded conditions, each defined by different combination of KCC2 and NKCC1 fractions (see **Figure 7** and **Figure 8**): 1) control condition (1.0 KCC2 and 1.0 NKCC1), 2) aPC-LBN males (0.80 KCC2 and 0.59 NKCC1), 3) aPC-LBN females (0.59 KCC2 and 0.48 NKCC1), 4) pPC-LBN males (1.45 KCC2 and 0.90 NKCC1), and 5) pPC-LBN females (0.34 KCC2 and 0.88 NKCC1). These conditions were used to simulate the hypothesis that deviations in KCC2 expression from control levels modulate neuronal activity via GABAergic synaptic transmission. Specifically, lower KCC2 levels were expected to enhance excitability due to chloride extrusion and more depolarizing GABAergic input, whereas higher KCC2 levels were expected to dampen activity. Similarly, increased NKCC1 levels were hypothesized to enhance excitability, while reduced NKCC1 expression would reduce it, due to their respective effects on intracellular chloride accumulation.

**Figure 7.**
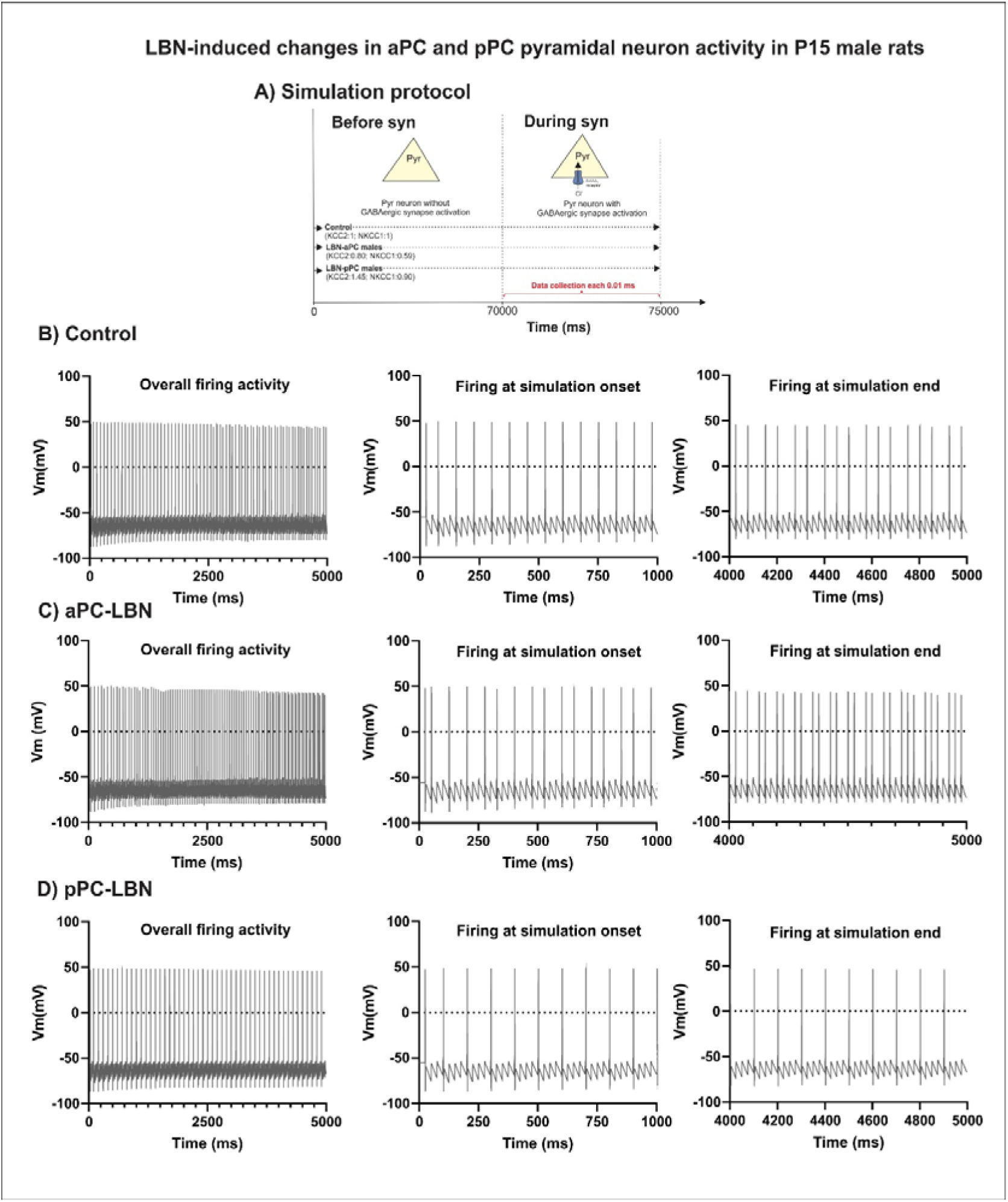
Dynamic changes of membrane potential activity of LBN-aPC and LBN- pPC neurons as a function of different fractions of KCC2/NKCC1 during GABAergic inputs in males. **A)** Experimental design of simulation before and after GABAergic synapse activation. **B**) spiking activity of the control P15 pyramidal neuron with a KCC2:NKCC1 fraction of 1:1 during a 5000 ms simulation, with GABAergic synaptic stimulation delivered every 20 ms. Firing activity was relatively stable, showing similar frequency during the first and last 1000 ms of the simulation. **C**) Spiking activity of an LBN-aPC neuron with a KCC2:NKCC1 fraction of 0.80:0.59. Firing was less frequent during the first 1000 ms and became more frequent during the last 1000 ms of the simulation compared to the control profile in B. **D**) Spiking activity of an LBN-pPC neuron with a KCC2:NKCC1 fraction of 1.45:0.90. Firing was consistently less frequent, both at the onset and the end of the simulation, compared to the control profile in A. The membrane potential was initialized at -55.54 mV, corresponding to the experimental resting potential of aPC pyramidal neurons at P14- P17. Data points were collected every 0.01 ms.

**Figure 8.**
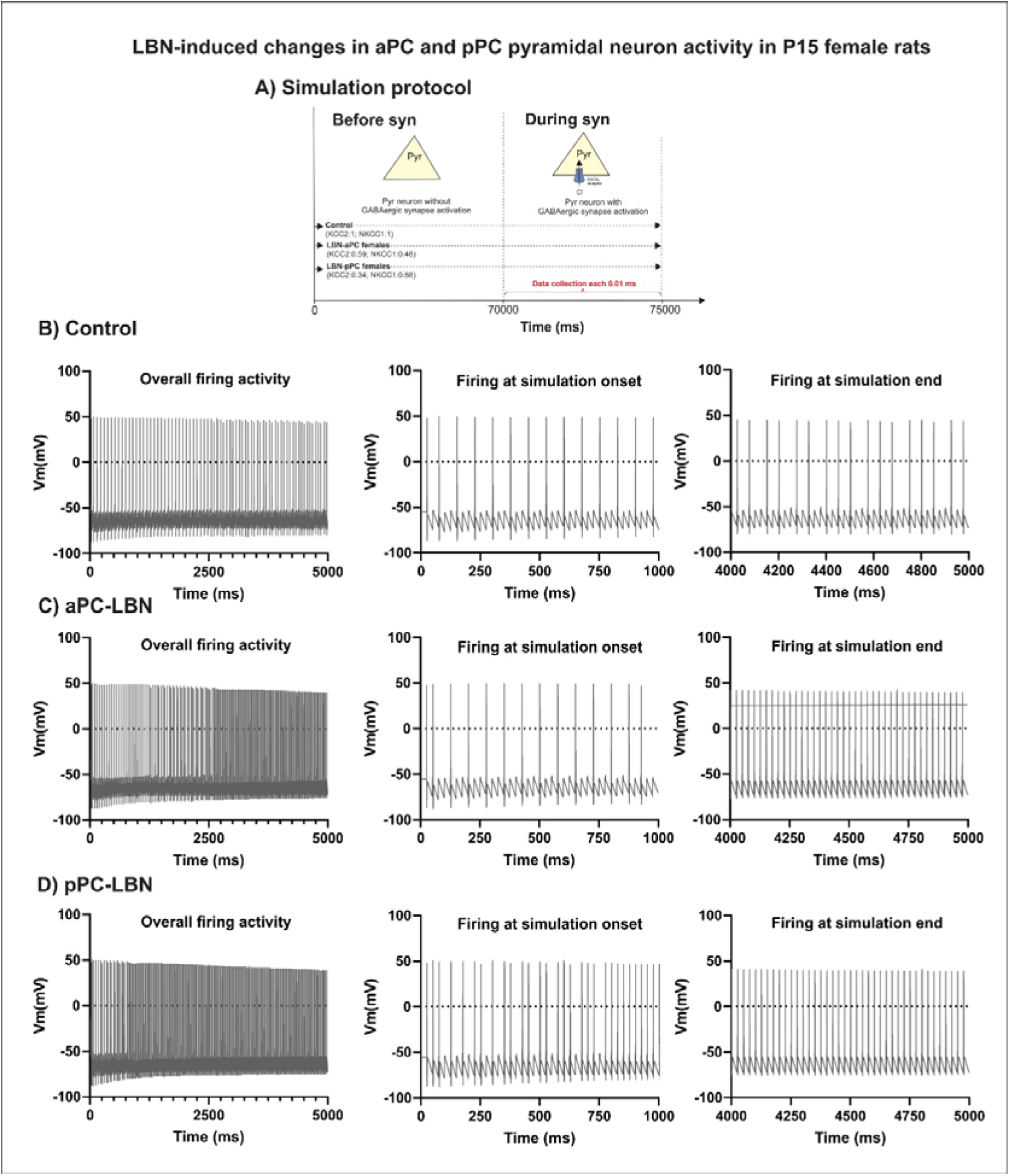
Dynamic changes of membrane potential activity of LBN-aPC and LBN- pPC neurons as a function of different fractions of KCC2/NKCC1 during GABAergic inputs in females. **A)** Experimental design of simulation before and after GABAergic synapse activation.**B**) spiking activity of the control P15 pyramidal neuron with a KCC2:NKCC1 fraction of 1:1 during 5000 ms simulation, with GABAergic synaptic stimulation delivered every 20 ms. Firing activity was relatively stable, showing similar frequency during the first and last 1000 ms of the simulation. **C**) Spiking activity of an LBN-aPC neuron with a KCC2:NKCC1 fraction of 0.59:0.48.

To explore these hypotheses, we conducted simulations totaling 750000 ms, with each condition simulated independently following an AB protocol (**Figure 7, A**). During the first phase (*Before syn*), the neuron was simulated for 70000 ms without GABAergic synaptic activation. Each neuron was initialized with its corresponding KCC2 and NKCC1 fractions and a membrane potential of -55.53 mV. Then, at 70000 ms, GABAergic synaptic transmission was continuously activated and maintained for 5000 ms (*During syn*). During this window, data were collected at 0.01 ms intervals to analyze the membrane potential, the number of spikes, intracellular chloride concentration, and chloride reversal potential. We present the results separately for male and female rats.

***aPC and pPC-LBN males*.** The simulation under control conditions, representing P15 animals, exhibited a constant spiking activity, with 14 spikes in the first 1000 ms and 17 spikes in the last 1000 ms of the simulation (**Figure 7,B**). A reduction of 20% KCC2 and 41% in NKCC1 (aPC-LBN male condition) also induced spiking activity, but with different temporal patterns. A total of 16 spikes were recorded during the first 1000 ms, and 27 spikes were recorded during the last 1000 ms, indicating an increase in firing frequency over time (**Figure 7,C**).

In contrast, the pPC-LBN male condition, characterized by a 45% increase in KCC2 and 10% decrease in NKCC1, also produced spiking activity, but with a more stable firing pattern across the simulation, with 11 spikes during the first 1000 ms and 9 spikes during the last 1000 ms of the simulation (**Figure 7,C).**

***aPC and pPC-LBN females*.** To compare female profiles, we used the same control applied to males (**Figure 8,B**). In the aPC-LBN female condition, characterized by a 41% reduction in KCC2 and a 52% reduction in NKCC1, the neuron showed regular spiking activity, with 14 spikes in the first 1000 ms, increasing frequency to 40 spikes in the last 1000 ms (**Figure 8, C).** Similarly, the pPC-LBN female condition, with a 66 % reduction in KCC2 and a 12% reduction in NKCC1, induced high-frequency firing. A total of 30 spikes were recorded in the first 1000 ms, and 40 spikes in the last 1000 ms simulation (**Figure 8, D).**

Spiking activity profiles for both male and female conditions are summarized in **Figure 9**. In males, the aPC-LBN condition exhibited increased spiking activity compared to the pPC-LBN condition, but lower than control levels, while pPC-LBN condition showed reduced spiking relative to control (**Figure 9, A**). In females, the aPC-LBN condition also displayed elevated spiking activity compared to control, whereas the pPC-LBN condition showed even higher activity than both control and aPC-LBN conditions (**Figure 9, B**).

**Figure 9.**
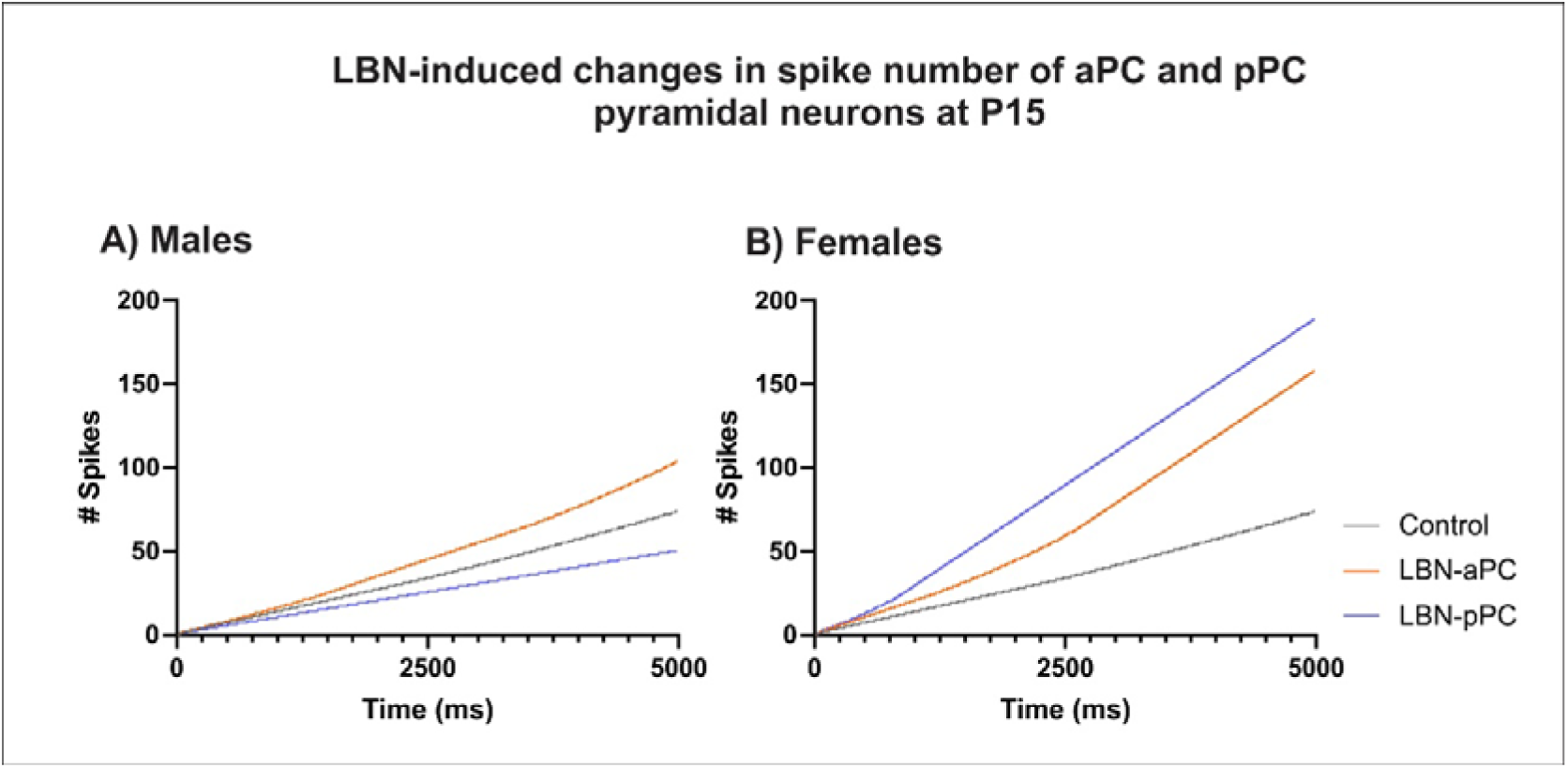
Summarized profile of spiking activity of P15 LBN-aPC and LBN-pPC neurons in males and females. **A)** Spiking activity profiles of control, LBN-aPC, and LBN-pPC neurons in males. The LBN-aPC profile showed increased firing frequency, while LBN-pPC profile showed decreased frequency compared to the control profile. **B**) The same control profile as in (A) was used to compare the profiles of LBN-aPC and LBN-pPC in females. Both LBN-aPC and LBN-pPC profiles showed increased firing compared to control, with LBN-pPC exhibiting the highest frequency, even greater than LBN-aPC.

Firing was more frequent during the first 1000 ms and became even more frequent during the last 1000 ms of the simulation compared to the control profile in (B). **D**) Spiking activity of a LBN-pPC neuron with KCC2:NKCC1 fraction of 0.34:0.88. Firing was markedly more frequent, both at the onset and the end of the simulation, compared to the control profile in (A). The membrane potential was initialized at -55.54 mV, corresponding to the experimental resting potential of aPC pyramidal neurons at P14- P17. Data points were collected every 0.01 ms.

The dynamic of intracellular chloride concentration [Cl^-^]_e_, chloride reversal potential (E_Cl_), and spike accumulation in response to GABAergic synaptic stimulation are shown in **Figure S7** (males) and **Figure S8** (females). Under control conditions, [Cl^-^]_e_ remained relatively stable during the initial period and increased slightly over time, accompanied by a mild depolarization shift in E_Cl_ and a linear accumulation of spikes (**Figure S7, A**). In the aPC-LBN male condition (**Figure S7, B**), [Cl^-^]_e_ showed a moderate increase over time, with a progressive depolarization of E_Cl_, consistent with increased firing activity. In contrast, the pPC-LBN male condition (**Figure S7, C**) exhibited a slight early reduction in [Cl^-^]_e_ , followed by stabilization, and E_Cl_ remained more hyperpolarized compared to the control and aPC-LBN conditions. The spike accumulation curve was also flatter, reflecting reduced firing. In females, both LBN conditions led to more pronounced changes in chloride homeostasis (**Figure S8**). The aPC-LBN female condition (**Figure S8, B**) exhibited a continuous increase in [Cl^-^]_e_ and robust depolarization of E_Cl_, in parallel with a higher spike accumulation rate than controls (**Figure S8, A)**. This trend was even more accentuated in the pPC-LBN female condition (**Figure S8, C**), where [Cl^-^]_e_ increased progressively and E_Cl_ shifted markedly towards depolarized values, supporting the strong increase in spike frequency observed in this condition.

Together, these results indicate that chloride dynamics and neural excitability under LBN are modulated by the specific combination of KCC2 and NKCC1 fractions modeled in each condition. The patterns of neural activity in each condition highlight how imbalances in KCC2 and NKCC1 fractions rather than absolute expression levels can critically shape GABAergic signaling outcomes in developing pyramidal neurons.

## 4. Discussion

This study aimed to characterize the developmental expression of chloride cotransporters in the piriform cortex during early postnatal development under typical conditions and to assess how variations in early maternal caregiving, induced by the LBN paradigm, influence this trajectory. We focused on KCC2 and NKCC1 because the piriform cortex is a region both necessary and sufficient for attachment learning in infant rats (Sullivan, 2003; Yuan *et al*., 2014). Previous work from our group has shown that, within this region, GABAergic synaptic signaling to layer 2/3 pyramidal cells undergoes marked maturation (Pardo, 2018; Pardo *et al*., 2018), in parallel with changes in passive and active intrinsic properties (Oruro *et al*., 2020a). During this stage, GABA signaling remains depolarizing and contributes to excitatory drive, conferring the olfactory bulb-piriform cortex circuit with exceptionally rapid odor-learning capabilities that characterize the sensitive period for attachment learning (Oruro *et al*., 2020b). The present results indicate that, as in other developing brain regions, the levels of KCC2 and NKCC1 increase progressively with age, reaching adult-like levels by weaning in both sexes. However, LBN exposure during the sensitive period reduced protein levels of both cotransporters in both sexes in the aPC at P15, while inducing opposite effects in the pPC. Computational modeling based on these profiles revealed distinct firing patterns in response to isolated GABAergic input, showing higher firing frequency when both KCC2 and NKCC1 were low, and elevated firing when KCC2 was low and NKCC1 was high, suggesting that developing GABAergic circuits in the piriform cortex are highly sensitive to maternal caregiving variations, with potential consequences for neuronal activity patterns.

### 4.1. Ontogenetic changes of KCC2 and NKCC1 expression in the piriform cortex

In several developing brain regions, KCC2 protein levels progressively increases with age, as reported in rodent cerebral cortex (Dzhala *et al*., 2005; Takayama & Inoue, 2010; Murguía-Castillo *et al*., 2013; Kovács *et al*., 2014; Markkanen *et al*., 2014; Zavalin *et al*., 2024), hippocampus (Rivera *et al*., 1999; Gulyás *et al*., 2001; Stein *et al*., 2004; Murguía-Castillo *et al*., 2013; Spoljaric *et al*., 2019), and olfactory bulb (Wang *et al*., 2005). In the piriform cortex, KCC2 transcripts are detectable from gestational day E15.5, coinciding with neuronal differentiation (Stein *et al*., 2004). Immunolabeling studies have also reported KCC2 protein already present at P0, initially confined to dendrites in the superficial layers at P0-P3, extending to deeper layers by P4, and by P6 showing strong labeling across all three layers, with a stable pattern through P13 (Kovács *et al*., 2014). In our study, KCC2 protein levels were low during the first postnatal week and progressively increased, reaching adult-like levels by weaning in both sexes.

For NKCC1, most cortical regions show high expression at birth, followed by a gradual decline (Marty *et al*., 2002; Ludwig *et al*., 2003; Dzhala *et al*., 2005). In contrast, our data reveal low NKCC1 expression levels in the piriform cortex during the first postnatal week, with a gradual increase thereafter, paralleling KCC2. KCC2 is neuron-specific, whereas NKCC1, though weakly expressed in neurons, is mostly expressed in non-neuronal cells (Russell, 2000; Kanaka *et al*., 2001; Stein *et al*., 2004; Uvarov *et al*., 2009). Thus, the gradual increase in both cotransporters may reflect the parallel maturation of neuronal and glial compartments.

The piriform cortex undergoes marked growth in thickness from birth to weaning, linked to neuronal differentiation (Sarma *et al*., 2011) dendritic maturation (Moreno-Velasquez *et al*., 2020), and synaptic remodeling (Westrum & Bakay, 1986; Westenbroek *et al*., 1988; Moriizumi *et al*., 1995; Franks & Isaacson, 2005; Poo & Isaacson, 2007; Sarma *et al*., 2011; Pardo *et al*., 2018; Martin-Lopez *et al*., 2019). At the microstructural level, KCC2 is predominantly localized to dendrites and plasma membrane domains by P12, with a substantial fraction also present in cytoplasmic vesicular compartments (Kovács *et al*., 2014), suggesting that the developmental increase reflects both dendritic expansion and intracellular accumulation. NKCC1 isoforms show cell-type specificity, with b isoform predominantly in neurons and a isoform predominantly in glial cells (Kurki *et al*., 2023), which could underline the parallel developmental trajectory observed here.

Sex differences in cotransporter expression have been described elsewhere: in the hypothalamus, males show higher levels of NKCC1 and lower KCC2 than females early postnatally (Perrot-Sinal *et al*., 2007); in the hippocampus and entorhinal cortex, NKCC1 protein rises in males but not females, whereas KCC2 increases earlier in females (Murguía-Castillo *et al*., 2013). In the piriform cortex, however, we found developmental upregulation of both cotransporters in both sexes at the protein level. NKCC1 mRNA mirrored protein in both sexes, but KCC2 mRNA showed a transient P11-P14 peak in females that was absent in males. This female-specific mRNA peak may relate to the postnatal rise in estradiol and gonadotropic hormones during this period (Döhler & Wuttke, 1975; Herath *et al*., 2001). NKCC1 is also sensitive to estrogens: in the hypothalamus, estradiol treatment does not alter phosphorylated NKCC1 in females but increases it in males, suggesting sex-dependent post-translational regulation (Perrot-Sinal *et al*., 2007). Moreover, sex differences may appear at the level of membrane localization without changes in total protein. In the hippocampus at P7, KCC2 membrane monomer levels are higher in females, a difference abolished by P19 and reduced by early testosterone treatment (Wolf *et al*., 2022), suggesting that gonadal hormones can modulate both cotransporters at multiple levels, a possibility that warrants direct investigation in the piriform cortex.

### 4.2. Altered maternal caregiving differentially shifts KCC2 and NKCC1 expression in anterior and posterior piriform cortex in a sex-dependent manner

A previous study using a similar LBN paradigm reported an accelerated GABA switch in the mPFC, driven by reduced NKCC1 mRNA levels at P9-P15 without altering KCC2 mRNA, consistent with reduced mEPSC frequency in mPFC pyramidal neurons (Karst *et al*., 2023). In contrast, our findings show that LBN exposure from P2-P9 resulted in region-, sex-, and transporter-specific alterations in the piriform cortex. In the aPC, LBN delayed the upregulation of both cotransporters, whereas in the pPC, it accelerated KCC2 expression while delaying NKCC1. These effects were further modulated by sex, with distinct expression profiles in males and females.

The LBN-induced shifts in cotransporter expression likely reflect the specific maternal caregiving profile in our cohort, characterized by a higher frequency of high-arched nursing posture and increased behavioral transition between nursing postures (Pardo *et al*., 2024). Such a caregiving pattern, beyond its known impact on stress-related systems in LBN paradigms (Walker *et al*., 2017), is likely to alter the temporal structure and quality of sensory cues reaching pups during the critical period of piriform maturation. These cues (particularly olfactory, tactile, and thermoregulatory) are essential for the experience-dependent maturation of developing circuits in various brain regions, as shown in studies of mother-infant interaction (Kojima *et al*., 2012; Kolb *et al*., 2017; Lapp *et al*., 2020; Nwabudike & Che, 2024; Nishizumi, 2025), and may be processed differentially in aPC and pPC. Given their distinct afferent profiles, these cues are likely processed differently in aPC and pPC, shaping neuronal activity, and consequently, the timing and extent of KCC2 and NKCC1 expression changes.

The distinct maturation profiles in aPC and pPC are likely linked to different afferent inputs and functional roles during early postnatal learning. The aPC receives direct input from LOT and associative projections from other cortical areas, whereas pPC integrates inputs from the aPC, the basolateral amygdala (BLA) and the LOT (Wilson and Sullivan 2011; Hagiwara et al. 2012; Luna and Morozov 2012). Functionally, the aPC is predominantly engaged during attachment learning, while pPC is more involved in aversion learning (Raineki *et al*., 2009). These differences in connectivity and function could drive region-specific patterns of neuronal activation and activity-dependent regulation of KCC2 and NKCC1 expression.

Sex further modulated these effects. In males, the LBN-induced acceleration of KCC2 expression was restricted to the pPC, whereas in females, changes were more uniform across regions. This divergence may reflect sex-biased maternal care, as males generally receive more licking than females (Moore & Morelli, 1979; Richmond & Sachs, 1984) potentially exposing each sex to qualitatively distinct sensory environments. Such differences in sensory input could contribute to the region-specific KCC2 expression profiles observed here.

### 4.3. Functional implications of LBN-induced changes in KCC2 and NKCC1 for membrane excitability of pyramidal cells during GABAergic signaling in the developing piriform cortex

High NKCC1 expression during the early postnatal period, together with reduced KCC2 levels, is known to underlie the initially excitatory action of GABA in several developing brain regions and the gradual upregulation of KCC2 drives the developmental switch in this GABA signaling from excitatory to inhibitory (Ben-Ari, 2002; Kaila *et al*., 2014), as E_GABA_ shifts from depolarizing to hyperpolarizing in parallel with the increase in KCC2 expression (Stein *et al*., 2004; Wang *et al*., 2005; Stil *et al*., 2009; Sun *et al*., 2013; Spoljaric *et al*., 2019; Raol *et al*., 2020). In our previous work, we showed that the GABA reversal potential shifts from more depolarized to more hyperpolarized values between <P10 and > P10 in rat pups (Pardo, 2018; Pardo *et al*., 2018). At <P10, E_GABA_ was even more depolarized than the resting membrane potential and threshold, progressively becoming hyperpolarized after P10 (Pardo 2018; Pardo et al. 2018; Oruro et al. 2020a; Oruro et al. 2020b).

Using these developmental dynamics as a framework, we implemented a biophysical model of the developing piriform cortex pyramidal neuron incorporating our experimental KCC2/NKCC1 protein levels data. The model captured the expected relationships in which lower KCC2 and higher NKCC1 proportions maintained a more depolarized E_GABA_, supporting greater GABA-driven excitability, whereas higher KCC2 expression shifted E_GABA_ towards hyperpolarization, reducing spiking.

Our results show that in the aPC of LBN pups at P15, delayed KCC2 upregulation maintained E_GABA_ depolarized (∼ -45 mV), increasing spiking during GABAergic input compared to control simulations. This effect was slightly more pronounced in females, consistent with their greater KCC2 reduction. In contrast, in the pPC of LBN males, accelerated KCC2 expression shifted E_GABA_ to less depolarized values (below -45 mV), markedly reducing spiking. In LBN females, delayed upregulation of both cotransporters maintained E_GABA_ depolarized (∼ -42 mV), supporting higher spiking than in their aPC.

The aPC is recruited during the sensitive period for attachment learning (Colombel *et al*., 2023), when excitatory synaptic plasticity between LOT terminals and aPC pyramidal cells is high (Poo & Isaacson, 2007) and GABA acts in a depolarizing manner (Oruro et al 2020b). After this period, the same conditioning for attachment no longer produces odor preference, synaptic plasticity decreases (Poo & Isaacson, 2007), and GABA signaling becomes progressively hyperpolarized (Pardo, 2018; Pardo *et al*., 2018). We can speculate that a delayed KCC2 maturation in the aPC could prolong the period during which GABAergic input facilitates attachment learning. In the pPC, which is co-activated with the basolateral amygdala (BLA) during odor-shock associative learning (Raineki *et al*., 2009)and receives glutamatergic projections from the BLA (Luna & Morozov, 2012; Sadrian & Wilson, 2015), reduced excitability might instead facilitate earlier engagement in aversive learning.

### 4.4. Limitations

NKCC1 and KCC2 must be integrated into the neuronal membrane to function as cotransporters (Russell, 2000; Zhang *et al*., 2006), yet our simulation relied on total protein levels, which may also include cytoplasmic transporters. This could limit the precision of the functional estimates, although the predicted effects on membrane potential are consistent with excitatory GABAergic action observed in previous work (Oruro et al. 2020a; Oruro et al. 2020b).

### 4.5. Conclusion

In the developing piriform cortex of typically reared rats, KCC2 expression increased with age, whereas NKCC1 also showed a gradual increase, with subtle sex-specific transcriptional differences. Early maternal caregiving alteration through the LBN paradigm disrupted developmental trajectories in P15 pups in a region- and sex-dependent manner, delaying both cotransporters upregulation in the aPC while accelerating KCC2 and delaying NKCC1 in the pPC. Our computational simulation, parameterized with these molecular data, suggests that such alterations may modify the effectiveness of GABAergic signaling and the excitability of pyramidal neurons during a critical window of circuit maturation. This integrative approach provides a mechanistic insight into how caregiving-related changes in chloride transporter regulation can extend from molecular alterations to synaptic function and cellular activity, with potential implications for attachment-related circuit formation and development.

## Authors contribution

**C.B**. designed and performed the wet experiments, acquired and analyzed data, and contributed to drafting the manuscript. **E.M.O**. developed the computational model, performed simulations, and interpreted results. **C.M.S**. contributed to the experimental design, performed experiments, and assisted with data interpretation. **G.E.P.** conceptualized the study, designed the experiments, supervised the research, analyzed data, interpreted results, performed data visualization, wrote the manuscript, and secured funding. **L.F.P.O**. contributed to the experimental design, provided laboratory resources, supervised wet experiments, assisted with data interpretation, and revised the manuscript. All authors contributed to revising the manuscript and approved the final version for submission.

## Acknowledgment

The authors want to thank Sergio Cruz-Visalaya, Lucero B. Cuevas, and Alondra Casas Pary for their assistance during the experiments.

