## Supplementary material for "Maternal care modulates chloride cotransporter development during inhibitory circuit maturation in the piriform cortex": Supplementary Material .pdf

### 1. Detailed Methods of Experiment 1

#### 1.1. Tissue Collection and Preparation Protocol

Whole piriform cortex tissue was collected at eight postnatal ages: P5-6, P7-8, P11-12, P13-14, P15-16, P19-20, P21-22, and P134. A total of 25 litters were used in this experiment. For each group, two pups per litter were randomly selected and euthanized in a separate room. Pups younger than PND 16 were euthanized by rapid decapitation with sharp scissors, whereas older animals were euthanized using a guillotine. Adult rats (PND 134) were first anesthetized with a ketamine/xylazine cocktail prior to decapitation. Brains were quickly removed and placed ventral side up on a Petri dish over crushed ice. The piriform cortex was dissected using curved fine forceps, with the optic chiasm and rhinal fissure as anatomical landmarks. Dissections of the piriform cortex were performed bilaterally in a caudo-rostral direction by gently pressing the forceps approximately 0-5-1 mm into the cortex. The dissected tissues were collected into pre-weighed 1.5 mL RNAase-free microtubes, reweighed, frozen on dry ice, and stored at -80°C until further processing.

#### 1.2. RNA Extraction Protocol

Total RNA was extracted from piriform cortex samples using a modified TRIzol protocol. Briefly, 30-50 mg of tissue was placed in 1.5 mL RNAse-free tubes on ice. Samples were lysed by adding 100 µL of Mammalian Lysis Buffer supplemented with 5% 2-mercaptoethanol, followed by homogenization with a sterile glass grinder. This step was repeated until a total of 300 µL of lysis buffer had been added. Subsequently, 500 µL of TRIzol™ reagent (Invitrogen, USA) was added, vortexed for 15 sec, and incubated for 5 min at 4°C. Phase separation was achieved by sequential addition of chloroform (2 x 200 µL), followed by centrifugation at 8855 x g for 15 min at 4°C after each addition. The aqueous phase was collected and mixed with 1 mL of isopropanol, incubated at 20°C for 30 min, and centrifuged at 8855 x g for 15 min at 4°C. The resulting RNA pellet was washed twice with 75% ethanol (1 mL each), with each wash followed by centrifugation at 3459 x g for 5 min at 4°C. The pellet was air-dried at room temperature and resuspended in 10-20 µL of RNAse-free water. RNA concentration and purity were determined by spectrophotometry (NanoDrop 2000, Thermo Scientific, USA), with acceptable 260/280 and 260/230 ratios above 1.8. RNA integrity was assessed by 1% agarose gel electrophoresis, verifying the presence of distinct 28S, 18S, and 5S rRNA bands. Residual genomic DNA was removed by DNase treatment using the TURBO DNA-free™ Kit (Invitrogen, USA). Purified RNA was stored at -80 °C until cDNA synthesis.

#### 1.3. Western Blot Protocol

Protein expression levels of KCC2 and NKCC1 in the piriform cortex were evaluated by Western blot analysis. Tissue samples were lysed in 100 µL Mammalian lysis buffer supplemented with a protease inhibitor cocktail (TRIS-HCl 25 mM pH 8, NaCl 100 mM, Triton X-100 0.1%, SDS 0.1%, EDTA 1 mM, TPCK 100 µM, TLCK 100 µM, AEBST 1 mM, Leupeptin 50 µM, E-G4 5 µM, Pepstatin A 20 µM). Homogenization was performed using a sterile glass grinder, followed by vortexing for 2 min. Lysates were incubated on crushed ice for 1 h and centrifuged at 8855 x g for 30 min at 4°C. The total protein supernatant was collected and stored at -20°C until analysis. Protein concentration was estimated using the Bradford method with BSA as a standard. For each sample, 50 µg of total protein was mixed with 6x loading buffer (TRIS-HCL 150 mM, pH 8.8, SDS 150 mM, bromophenol blue 0.04 g, glycerol 20%, 2-mercaptoethanol 10%) and denatured at 95 °C for 5 min. Samples were cooled to room temperature before loading onto SDS-polyacrylamide gels.

Electrophoresis was performed at constant voltage (starting at 100 V and increasing to 150 V) until complete migration of the loading dye. Proteins were transferred onto PVDF membranes using a semi-dry transfer system at 400 mA for 1 h in transfer buffer (25 mM Tris, 192 mM glycine, 5% methanol, pH 8.3). Membranes were then blocked for 1 h at room temperature in 5% nonfat dry milk in PBST (0.1 % Tween-20). Membranes were incubated overnight at 4 °C with the following primary antibodies diluted in 1% milk/PBST: rabbit anti-actin (1:4000, Thermo Fisher, MA5-32479), rabbit anti-KCC2 (1:2000, Thermo Fisher, PA5-110374), and rabbit anti-NKCC1 (1:2000, Invitrogen, PA5-95145). After three washes in PBST (10 minutes each), membranes were incubated for 1 hour at room temperature with a horseradish peroxidase (HRP)-conjugated biotinylated anti-rabbit secondary antibody (1:5000, Invitrogen, 656140). Subsequently, membranes were incubated for 1 h in ABC reagent (Vectastain ABC Kit, Vector Laboratories), and immunoreactive bands were visualized using DAB substrate (DAB Peroxidase Substrate Kit, Vector Laboratories). Membranes were imaged under uniform light exposure, and band intensities were quantified using ImageJ software.

Protein expression data of KCC2 and NKCC1 were normalized to β-actin to control loading variability and then expressed as percentages relative to the mean expression level observed in adult rats (P134) within the same experiment. This normalization enabled comparative analysis across early developmental time points (P5–8, P11–14, P15–18, and P19–22).

#### 1.4. Quantitative RT-PCR Protocol

cDNA was synthesized from 500 ng of purified total RNA using a conventional endpoint reverse transcription PCR protocol. The reaction included two master mixes in equal proportions (1:1). Master Mix 1 contained 2.0 µL of 10x RT buffer, 0.8 µL of 25x dNTPs, 2.0 µL of 10x random primers, 1.0 µL of reverse transcriptase, and 4.2 µL of DEPC-treated water. Master Mix 2 consisted of the RNA sample diluted in DEPC-treated water to reach the desired concentration. The combined mixes were incubated in a thermocycler at 25 °C for 10 minutes, 37 °C for 120 minutes, 85 °C for 5 minutes, and finally held at 4 °C.

Quantitative PCR was then performed using SYBR Green detection chemistry and the BlasTaq™ 2X qPCR MasterMix (Applied Biological Materials Inc., Canada), in a final reaction volume of 10 µL. Each reaction consisted of 5 µL of 2X SYBR Green MasterMix, 1 µL of forward/reverse primers (2 µM), 2 µL of cDNA (50 ng/µL), and 2 µL of RNAse-free water. Additionally, 1 µL of ROX reference dye was added per 1.25 mL of master mix, as recommended by the manufacturer. Reactions were conducted in an AriaMX Real-Time PCR System (Agilent, USA), equipped with FAM/SYBR optical modules. Thermal cycling conditions consisted of an initial denaturation at 95 °C for 3 minutes, followed by 40 amplification cycles of denaturation at 95 °C for 10 seconds and annealing/extension at 60 °C for 15 seconds. Melting curve analysis was subsequently performed at 95 °C, 65 °C, and 95 °C, each for 30 seconds, to confirm single-product amplification and assess specificity. To validate amplification efficiency and linearity, standard curves were generated using five serial dilutions of cDNA from adult rat hippocampus (known to express KCC2 and NKCC1) at final concentrations of 100, $10^{-1}$, $10^{-2}$, $10^{-3}$, and $10^{-4}$ ng/µL. Each dilution was prepared by transferring 1 µL of the previous dilution into 9 µL of RNAse-free water. Three replicate reactions were run per dilution for each gene (gapdh, hprt1, kcc2, nkcc1). Amplification efficiencies calculated from the standard curves were 83.5% for GAPDH, 98.8% for Hprt1, 79.4% for KCC2, and 97.3% for NKCC1. Gene-specific primers were used to amplify KCC2 (, NKCC1, GAPDH, and

Hprt1. Amplicon sizes were 101 bp (KCC2), 120 bp (NKCC1), 142 bp (GAPDH), and 141 bp (Hprt1). Primer sequences are detailed in **Table S1**:

**Table S1. Sequence of primers, amplicon sizes, and efficiency**

| Gene | Sequence | GenBank Accession number | Amplicon size (bp) | Efficiency (%) |
|---|---|---|---|---|
| KCC2 | Forward: 5'- ATTGTGGGCTCTGTCCTTGA-3'<br>Reverse 5'- TTCACCTTCTCAGCCTCCAT-3' | NM_134363.2 | 101 | 79.4% |
| NKCC1 | Forward 5'- AACACACTTGTCCTCGGATTT-3'<br>Reverse 5'- GCGAATGACCACAACTCCAT-3' | NM_031798.2 | 120 | 97.3% |
| GAPDH | Forward 5'- CATCTTCCAGGAGCGAGATC-3'<br>Reverse 5'- GGAGATGATGACCCTTTTGG-3' | NM_017008 | 142 | 83.5% |
| Hprt1 | Forward 5'- GGACCTCTCGAAGTGTTGGA-3'<br>Reverse 5'- CTTTTCCACTTTCGCTGATGA-3' | NM_012583.2 | 141 | 98.8% |

Experimental quantification included 25 cDNA samples from five developmental age groups: P 5–8, P 11–14, P 15–18, P 19–22, and adult controls (P 134), with five biological replicates per group. For each qPCR run (96-well format), one sample from each group was randomly selected and analyzed in triplicate for both target genes (KCC2, NKCC1) and reference genes (GAPDH, Hprt1). RNAse-free water was included as a no-template control. Fluorescence data were analyzed using AriaMX software (Agilent, USA), which provided quantification cycle (Cq) values, normalized fluorescence (ΔRn), and melting temperatures (Tm). Only samples with Cq values within ±1 of the group mean, Tm within ±0.25 °C, and no evidence of primer dimer formation were included in the final analysis. Relative expression levels were calculated using the ΔΔCq method, normalized to the geometric mean of GAPDH and Hprt1 (Taylor et al. 2019). Primer information is provided in **Table S1**. Amplification efficiencies were confirmed with standard curves from hippocampal cDNA.

All mRNA expression levels were expressed as a percentage relative to the mean expression level of adult rats (P134) from the same experiment, providing a developmental comparison across postnatal time points (P5–8, P11–14, P15–18, and P19–22).

#### 1.5. Immunochemistry Protocol

Three naïve animals per sex were deeply anesthetized (ketamine 1.0 µL/g, Xylazine, 0.5 µL/g) and transcardially perfused with 0.9% saline, followed by 4% paraformaldehyde (PFA) containing 10% picric acid (Sigma, 36011) in PB. Brains were post-fixed in PFA for 24 h at 4°C and subsequently cryoprotected in 30% sucrose until they sank. Coronal sections (50 µm thick) were obtained using a cryostat (CM 1520, Leica, Germany) at anatomical levels corresponding to the aPC and pPC, based on a developmental rat brain atlas (Khazipov et al. 2015). Sections were collected into 24-well plates containing 1X PBS and stored at 4°C.

For immunostaining, free-floating sections were washed three times for 5 min each in PBS 1X with 0.01% Tween-20 (PBST) under gentle orbital agitation (50 rpm). Non-specific binding was blocked by incubating sections in PBST containing 5% normal goat serum for 10 min with no agitation, followed by 1 h at 30 rpm gentle agitation. Sections were then incubated overnight at 4°C with primary antibodies diluted in PBST with 2.5% normal goat serum. The following primary antibodies were used: rabbit anti-KCC2 (1:1000, PA5-1010374, Thermo), rabbit anti-NKCC1 (1:1000, PA5-95145, Invitrogen), mouse anti-NeuN (1:1000, 266011, Synaptic Systems). After three additional 5 min washes in PBST, sections were incubated for 1 h at room temperature with fluorophore-conjugated secondary antibodies diluted in PBST with 2.5% normal goat serum. The following secondary antibodies were used: donkey anti-rabbit Alexa Fluor 647 (1:500, 711-605-152, Jackson ImmunoResearch), horse anti-mouse DyLight 594 (1:500, DI-2594, Vector Laboratories). From this point forward, all procedures were conducted in the dark to protect fluorophores from degradation. After final washes, sections were mounted on gelatin-coated glass slides. Tissue was carefully positioned using a fine brush, and excess liquid was removed from the margins. Slides were coverslipped with Fluoromount-G mounting medium (SouthernBiotech) and allowed to dry overnight at room temperature. Fluorescence imaging was performed the following day using a Zeiss Axio Observer 5 inverted microscope equipped with an Apotome 3 optical sectioning system, using 10x and 40x objectives.

This qualitative approach allowed confirmation of neuronal expression patterns without introducing bias from fluorescence intensity quantifications.

### 2. Detailed Method of Computational Modeling

#### 2.1. Overview of the Model

We implemented a conductance-based Hodgkin-Huxley (HH) type model incorporating electrodiffusive ion dynamics to simulate the activity of a pyramidal neuron in the anterior piriform cortex (aPC) (**Figure S1A**). The model was adapted from Lemaire et al (2021), who originally developed a two-neuron model (a pyramidal and a GABAergic neuron) to study the effect of sodium channel mutations in hippocampal excitability and epileptogenesis.

For our purposes, we focused exclusively on the pyramidal neuron component, as our central hypothesis concerned the role of KCC2 and NKCC1 in shaping GABAergic effects on piriform cortex pyramidal neurons during early postnatal development. We excluded the GABAergic neuron and all excitatory glutamatergic synapses, retaining only the postsynaptic effect of GABAergic input via GABAA receptor activation.

The pyramidal neuron model includes (**Figure S1B)**:

- Voltage-gated sodium ($Na^+$), potassium ($K^+$), and calcium ($Ca^{2+}$) channels
- Passive leak currents for $Na^+$, $K^+$, and $Cl^-$.
- $Na^+$/$K^+$-ATPase pump
- KCC2 and NKCC1 chloride cotransporters
- Intracellular and extracellular ion dynamics governed by mass conservation
- GABAergic input modeled as a direct synaptic activation, bypassing presynaptic dynamics.

Our implementation retained the full set of differential equations and parameters for the pyramidal neuron described by Lemaire et al (2021), with modifications introduced to simulate the ionic environment and transporter profiles measured experimentally at P15. These modifications allowed us to simulate chloride homeostasis and the impact of transporter expression on GABAergiç signaling during a critical window of cortical development.

#### 2.2. Model Implementation and Numerical Integration

The complete set of equations for the pyramidal neuron model was implemented in NetLogo, a multi-agent modeling environment, using an explicit Euler method with a fixed time step of 0.01 ms, different from the original model by Lemaire et al (2021), which used a 4th-order Runge-Kutta method in XPPAUT.

All variables and equations related to the GABAergic neuron were excluded. We also omitted excitatory glutamatergic synapses. The model instead simulates postsynaptic $GABA_A$ receptor activation as a sole source of synaptic input, allowing us to isolate the contribution of chloride transporters to GABAergic effects.

The model implementation included:

- Differential equations describing membrane voltage, ion concentrations, gating variables, pump and transporter activity, and synaptic state.
- Mass-conserving ion dynamics, with intracellular and extracellular concentrations updated at each time step, allowing for time-varying Nernst potentials.
- Simplified synaptic input, with GABAergic activations represented as a direct step input to the chloride conductance.

The resting membrane potential ($V_{rest}$) was manually adjusted to -55.54 mV, consistent with experimental measurements from pyramidal neurons in the aPC at P14-P17 (Oruro et al. 2020).

### 2.3. **Model Components and Equations**

#### 2.3.1. **Membrane Potential Dynamics**

The membrane potential dynamics of the pyramidal neuron were modeled using the classical Hodgkin-Huxley (HH) framework. The change in membrane voltage over time is determined by the sum of ionic currents crossing the membrane, which include active (voltage-gated), passive (leak), synaptic, cotransporter, and pump components. The equation governing membrane voltage is:

$$C \frac{dV_e}{dt} = -I_{Na,e} - I_{K,e} - I_{Cl,e} \quad \text{(Equation 1)}$$

where:

- $C$ is the membrane capacitance,
- $I_{Na,e}$, $I_{K,e}$, and $I_{Cl,e}$, are the total sodium, potassium, and chloride currents across the membrane, respectively.

Each ionic current includes contributions from voltage-gated channels, leak components, transporters (KCC2, NKCC1), and pump ($Na^+/K^+$-ATPase). The relative balance of these currents determines whether the neuron depolarizes or hyperpolarizes at the given moment in time. The model was initialized and tuned to ensure convergence to this resting potential under baseline conditions, as further described in **Section 2.3.7**.

#### 2.3.2. **Gating Variables**

The dynamics of sodium and potassium channels were modeled using standard HH gating variables, which regulate the opening and closing of ion channels as functions of voltage. Three gating variables were used:

- $m_e$, sodium activation
- $h_e$, sodium inactivation
- $n_e$, potassium activation

Their time evolution is governed by first-order kinetics:

$$\frac{dm_e}{dt} = \alpha_{m,e}(1 - m_e) - \beta_{m,e}.m_e \qquad \text{(Equation 2)}$$

$$\frac{dh_e}{dt} = \alpha_{h,e}(1 - h_e) - \beta_{h,e}.h_e \qquad \text{(Equation 3)}$$

$$\frac{dn_e}{dt} = \alpha_{n,e}(1 - n_e) - \beta_{n,e}.n_e \qquad \text{(Equation 4)}$$

The voltage-dependent rate constants (α and β ) were adapted from Lemaire et al (2021), with the modification of the sodium activation function ($\alpha_{m,e}$), which was adjusted from -54 mV to -48 mV to increase the action potential threshold and match the target resting potential of -55.54 mV. The rate functions are:

Originally:

$$\alpha_{m,e} = 0.32 \frac{V_e + 54}{1 - \exp\left(-\frac{V_e + 54}{4}\right)}$$

Adjusted to:

$$\alpha_{m,e} = 0.32 \frac{V_e + 48}{1 - \exp\left(-\frac{V_e + 48}{4}\right)}$$

$$\beta_{m,e} = 0.28 \frac{V_e + 27}{\exp\left(\frac{V_e + 27}{5}\right) - 1},$$

$$\alpha_{h,e} = 0.128 \exp\left(-\frac{V_e + 50}{18}\right),$$

$$\beta_{h,e} = \frac{4}{1 + \exp\left(-\frac{V_e + 27}{5}\right)},$$

$$\alpha_{n,e} = 0.032 \frac{V_e + 52}{1 + \exp\left(-\frac{V_e + 52}{5}\right)},$$

$$\beta_{n,e} = 0.5\exp\left(-\frac{V_e + 57}{40}\right)$$

The steady-state calcium activation was defined as:

$$m_{\infty,Ca} = \frac{1}{1 + \exp\left(-\frac{V_e + 25}{25}\right)}$$

These gating variables dynamically modulate the conductance of fast sodium and delayed-rectifier potassium channels, thereby shaping action potential generation and repolarization.

#### 2.3.3. Intracellular Ion Dynamics

The intracellular concentrations of potassium, sodium, chloride, and calcium ions were dynamically updated at each time of the simulation. These concentrations evolved based on the net transmembrane currents for each ion, scaled by the conversion factor $\gamma_e$, which translates current ($\mu A/cm^2$) into changes in concentration (mM/ms) based on geometric properties of the modeled cell.

The conversion factor $\gamma_e$ was originally calculated by Lemaire et al (2021) based on the assumption of a spherical geometry for both pyramidal and GABAergic neuron. In their model, the GABAergic neuron had a volume equal to 2/3 of the pyramidal neuron. In our implementation, we excluded the GABAergic neuron and recalculated the conversion factor $\gamma_e$ specifically for the pyramidal neuron and its corresponding extracellular space. The moment-to-moment changes in intracellular concentration were determined by the following equations:

$$\frac{d[K^+]_e}{dt} = -\gamma_e . I_{K,e} \quad \text{(Equation 5)}$$

$$\frac{d[Na^+]_e}{dt} = -\gamma_e . I_{Na,e} \quad \text{(Equation 6)}$$

$$\frac{d[Cl^-]_e}{dt} = -\gamma_e . I_{Cl,e} \quad \text{(Equation 7)}$$

$$\frac{d[Ca^{+2}]_e}{dt} = -\frac{\gamma_e}{2} . I_{Ca,e} - \frac{[\ Ca^{2+}]_e}{\tau_{Ca}} \quad \text{(Equation 8)}$$

#### 2.3.4. Extracellular Ion Dynamics

In our model, only pyramidal neurons were simulated. Therefore, the contribution of GABAergic neurons and their associated extracellular compartments was excluded. Extracellular ion concentrations were updated based on the following equations:

$$\frac{d[K^+]_o}{dt} = \frac{Vol_e}{Vol_o} . \gamma_e . I_{k,e} - I_{k,diff} \quad \text{(Equation 9)}$$

$$\frac{d[Na^+]_o}{dt} = \frac{Vol_e}{Vol_o} . \gamma_e . I_{Na,e} \quad \text{(Equation 10)}$$

$$\frac{d[Cl^-]_o}{dt} = \frac{Vol_e}{Vol_o} . \gamma_e . I_{Cl,e} \quad \text{(Equation 11)}$$

Where:

- $Vol_e$ is the intracellular volume of the pyramidal neuron.
- $Vol_o$ is the perisynaptic extracellular volume.
- $\gamma_e$ is the conversion factor from ionic current to concentration.

To prevent non-physiological accumulation of potassium in the extracellular space during simulations, we implemented a diffusion term modeling the passive leakage of extracellular potassium into a bath:

$$I_{K,diff} = \varepsilon \left([K^+]_o - K_{bath}\right)$$

Where:

- $\varepsilon$ is the diffusion rate constant.
- $K_{bath}$ is the potassium concentration in the external bath

#### 2.3.5. GABAergic Synaptic Input

In this model, the GABAergic synaptic input to the pyramidal neuron was simplified. Since we exclude both the GABAergic neuron present in the original model by Lemaire et al (2021) model, the conditional dependence of synaptic activation on the membrane potential of the inhibitory neuron was removed. Instead, the inhibitory synapse was modeled as an external input such that the synaptic variable *Si* was set to 1 when GABAergic stimulation was simulated. This directly represents the activation (opening) of postsynaptic GABAA receptors mediating chloride influx, which was governed by the equation:

$$\frac{ds_i}{dt} = -\frac{1}{\tau_i} s_i \quad \text{(Equation 12)}$$

with the condition:

- $s_i \longleftarrow 1$ when GABAergic input is simulated

This variable *Si* directly modulates the GABAergic conductance $g_{GABA,e}$ and therefore the inhibitory current computed as:

$$I_{GABA,e} = g_{GABA,e} S_i \left(V_e - E_{Cl,e}\right)$$

#### 2.3.6. Ion currents and membrane potential dynamics Potentials

Ion currents were computed using reversal potential dynamically derived from Nerst Equation (Lemaire et al 2021), depending on intra-and extracellular ion concentrations at each time step:

$$E_{ion} = \frac{RT}{\mathrm{Z_{ion}}F} . \log\left(\frac{[\mathrm{ion}]_{\mathrm{extracellular}}}{[\mathrm{ion}]_{\mathrm{intracellular}}}\right)$$

where

- R=8,314 mJ.(k.mol)$^{-1}$t, he ideal gas constant
- T: temperature in Kelvin
- F= NA. $e$ , Faraday constant
- $Z_{ion}$: the valence of the ion

**Sodium currents**

- Fast inward currents (transient):

$$I_{Na,FI,e} = g_{Na,FI,e}\, m_e^3\, h_e \left(V_e - E_{Na,e}\right)$$

- Leak sodium current:

$$I_{Na,L,e} = g_{Na,L,e}(V_e - E_{Na,e})$$

**Potassium currents**

- Delayed rectifier potassium current:

$$I_{K,DR,e} = g_{K,DR,e}\, n_e^4\, h_e \left(V_e - E_{K,e}\right)$$

- Calcium-activated potassium currents:

$$I_{K,AHP,e} = g_{K,AHP,e} \frac{[Ca^{2+}]_e}{[Ca^{2+}]_e + K_{Ca}} \left(V_e - E_{K,e}\right)$$

- Leak potassium current:

$$I_{K,L,e} = g_{K,L,e}(V_e - E_{K,e})$$

**Chloride currents**

- Leak chloride current:

$$I_{Cl,L,e} = g_{Cl,L,e}(V_e - E_{Cl,e})$$

#### 2.3.7. Na+/K+ ATPase Pump

The $Na^+/K^+$ ATPase current plays a key role in maintaining ionic gradients and stabilizing the resting membrane potential. In this model, pump activity is regulated by both intracellular and extracellular ion concentrations and is modulated by membrane voltage. To reflect the experimental conditions of pyramidal neurons in aPC at P14-P17, model parameters were adjusted so that the membrane potential stabilizes at -55.54 mV under resting conditions.

The $Na^+/K^+$ pump current is defined as:

$$I_{pump,e} = \rho_{pump}\ (\ v_e)\ \left(\frac{[Na^+]_e}{[Na^+]_e + K_{pump,Na}}\right)^3 \left(\frac{[Na^+]_o}{[K^+]_o + K_{pump,k}}\right)^2$$

Where:

- $\rho_{pump}\ (\ v_e)$, maximum pump rate as a function of membrane voltage
- $K_{pump,Na}$, half-saturation constant for intracellular sodium
- $K_{pump,k}$ is the half-saturation constant for extracellular potassium
- $v_e$, membrane potential

The voltage dependence of the pump rate is modeled as: .

$$\rho_{pump}\ (v) =\ \rho_{pump,-55.54} \frac{f(v)}{f(-55.54)}$$

Where

$$f(\ v) = \frac{1 + \tanh\left(a\frac{F}{RT}v + b\right)}{2}$$

#### 2.3.8. Transmembrane Ion Cotransporters (KCC2 and NKCC1)

The potassium-chloride cotransporter (KCC2) and the sodium-potassium-chloride cotransporter (NKCC1) are essential components for regulating intracellular chloride levels and shaping GABAergic signaling during early development.

The KCC2 maintains low intracellular chloride concentration by extruding potassium and chloride using the potassium gradient:

$$I_{KCC} = \frac{\rho kcc}{\gamma_e} log\left(\frac{[K^+]_e[Cl^-]_e}{[K^+]_o[Cl^-]_o}\right)$$

The NKCC1 transporters sodium, potassium, and chloride into the cell (1Na:1K:2Cl):

$$I_{NKCC} = \frac{\rho NKCC}{1 + exp(K_{NKCC,K}[K^+]_o)} \left( \log\left(\frac{[K^+]_e[Cl^-]_e}{[K^+]_o[Cl^-]_o}\right) + \log\left(\frac{[Na^+]_e[Cl^-]_e}{[Na^+]_o[Cl^-]_o}\right)\right)$$

Where:

- $\rho KCC$ and $\rho NKCC$, maximal strengths (activity rates) of KCC2 and NKCC1
- $\gamma_e$, current-to-concentration conversion factor
- $K_{NKCC,K}$, half-activation constant of NKCC1 as a function of extracellular potassium concentration
- Subscripts e and o refer to intracellular and extracellular compartments, respectively.

These formulations, taken from Lemaire et al (2021), capture the dependence of cotransporter fluxes on the ionic gradients.

##### 2.3.9. Net Ionic Currents

At each simulation time step, the total transmembrane current for each ion was computed as the sum of its individual contributions from ion channels, cotransporters, and pumps. These cumulative currents drive the changes in membrane potential and ion concentrations described in previous sections.

The total sodium current was defined as:

$$I_{Na,e} = I_{Na,FI,e} + I_{Na,L,e} + 3I_{pump,e} + I_{NKCC}$$

The total potassium current was defined as:

$$I_{K,e} = I_{K,DR,e} + I_{K,AHP,e} + I_{K,L,e} + I_{KCC} + I_{NKCC} - 2I_{pump,e}$$

The total chloride current was defined as:

$$I_{Cl,e} = I_{Cl,L,e} - I_{KCC} - 2I_{NKCC} + I_{GABA,e}$$

##### 2.3.10. Conversion factors

To preserve mass conservation and properly ionic currents to changes in intra-and extracellular concentrations, we adopted and adapted the conversion originally described by Lemaire et al 2021.

- $\gamma_e$,current-to -concentration for intracellular space:

$$\gamma_e = \frac{S_e}{10^3 Vol_e N_A e}$$

Assuming spherical shape:

$$S_e = 4\pi \left(\frac{3Vol_e}{4\pi}\right)^{\frac{2}{3}}$$

Substituting values, we obtained:

$$\gamma_e = 4.45 * 10^{-5} \mathrm{mol.cm^2.\mu C^{-1}.L^{-1}}$$

- $\beta_1$, volume ratio of intracellular to extracellular space:

$$\beta_1 = \frac{Vol_e}{Vol_o} = 4$$

- $\beta_2$, scaling factor for extracellular volume:

$$\frac{Vol_e}{Vol_o} = \frac{\beta_1}{1 + \beta_1}$$

To enforce mass conservation (except for K+, which includes extracellular diffusion), we defined toral conserved quantities as:

- Sodium:

$$\mathrm{Na}_\Sigma = [Na^+]_o + \frac{Vol_e}{Vol_o}[Na^+]_e = 145 + 4 * 10 = 185 \text{ mM}$$

$$[Na^+]_o = \mathrm{Na}_\Sigma - \frac{Vol_e}{Vol_o}[Na^+]_e$$

- Chloride:

$$\mathrm{Cl}_\Sigma = -[Cl^-]_o \frac{Vol_e}{Vol_o}[Cl^-]_e = 130 + \frac{4}{1+\beta_1} * 5 = 142 \text{ mM}$$

$$[Cl^-]_o = \mathrm{Cl}_\Sigma - \frac{Vol_e}{Vol_o}[Cl^-]_e$$

Based on Lemaire et al (2021), we implemented the electrodiffusive constrain that links charges in membrane potential $V_e$ to the sum of ionic concentration changes:

$$\mathrm{C.}\frac{dV_e}{dt} - \frac{1}{\gamma_e}\left(\frac{d[Na^+]_e}{dt} + \frac{d[K^+]_e}{dt} - \frac{d[Cl^-]_e}{dt}\right) = 0$$

After removing the time derivatives and isolating the membrane potential, we derived the steady-state constraint:

$$\mathrm{H}_1 = Cv_e - \frac{1}{\gamma_e}([Na^+]_e + [K^+]_e - [Cl^-]_e)$$

Substituting baseline concentrations ($[Na^+]_e$ =10 mM, $[K^+]_e$= 140 mM, $[Cl^-]_e$ = 5 mM) and $V_e$=-55.54 mV, we obtained:

$$\mathrm{H}_1 = 70 - \frac{1}{\gamma_e}(10 + 140 - 5) \approx -3{,}258{,}497 \mu\mathrm{A.cm}^{-2}$$

Finally, potassium intracellular concentration can be expressed as a function of other variables:

$$[K^+]_e = \gamma_e\,(v_e - \mathrm{H}_1) - [Na^+]_e + [Cl^-]_e$$

#### 2.4. Initialization and Steady-Stare equilibration

Given the modifications introduced to the original model by Lemaire et al. (2021), including the exclusion of the GABAergic neuron and adjustments to transporter parameters, we initially observed large oscillations in the simulated variables. To ensure stability, we implemented the following approach:

a) The model was first run until all variables reached a dynamic steady state, defined by the cessation of oscillatory transients and convergence toward constant values over time.
b) Once steady-state values were achieved, we extracted the corresponding values of ionic concentrations and membrane potential and used them to reset the model´s initial conditions. This ensured that all subsequent simulations began from a stable baseline.

This approach allowed us to simulate a system operating around a realist equilibrium, rather than one distorted by numerical instability. The resulting resting membrane potential closely matched the experimental observed value for anterior piriform cortex pyramidal neurons at P15, which is -55.54 mV. This equilibrium served as the starting point for all hypothesis-driven simulations conducted in this study.

**Table S2. Parameters used in the conductance-based model of aPC pyramidal neuron**

| Description | Parameter | Value | Unit |
|---|---|---|---|
| **State variables (dynamics)** | | | |
| **Pyramidal neuron** | | | |
| Membrane potential | $v_e$ | - | mV |
| Sodium activated gating | $m_e$ | dynamic* | - |
| Sodium inactivated gating | $h_e$ | dynamic* | - |

| | | | |
|---|---|---|---|
| Potassium activating gating | $n_e$ | dynamic* | - |
| Intracellular potassium concentration | $[K^+]_e$ | dynamic* | mM |
| Intracellular sodium concentration | $[Na^+]_e$ | dynamic* | mM |
| Intracellular chloride concentration | $[Cl^-]_e$ | dynamic* | mM |
| Intracellular calcium concentration | $[Ca^{2+}]_e$ | dynamic* | mM |
| GABAergic synapse variable | $S_i$ | dynamic (0-1)* | - |
| **Extracellular concentrations** | | | |
| Extracellular potassium concentration | $[K^+]_o$ | dynamic* | mM |
| Extracellular sodium concentration | $[Na^+]_o$ | dynamic* | mM |
| Extracellular chloride concentration | $[Cl^-]_o$ | dynamic* | mM |
| Ions reversal potentials | $E_{Na}, E_K, E_{Cl},$ | dynamic* | mM |
| **Model parameters (fixed)** | | | |
| **General** | | | |
| Time | $t$ | 0.01* | ms |
| Elementary charge | $e$ | $1.6 \times 10^{-19}$ | C |
| Avogadro number (Na) | | $6.02 \times 10^{23}$ | $mol^{-1}$ |
| Ideal gas constant | R | 8.314 | $mJ.K^{-1}.mol^{-1}$ |
| Temperature | T | 310 | K |
| Membrane capacitance per area unit | $C$ | 1* | $\mu F.cm^{-2}$ |
| Pyramidal neuron volume | $Vol_e$ | $1.4368 \times 10^{-9}$ * | $cm^3$ |
| Temperature | $T$ | 309.15* | K |
| Surface area | $S_2$ | - | |
| Pump maximal rate at −55 mV | $\rho_{pump, -55}$ | 27 * | $\mu A.cm^{-2}$ |
| Pump half activation intracellular $[Na^+]$ | $K_{pump, Na}$ | 7.7 * | mM |
| Pump half activation extracellular $[K^+]$ | $K_{pump, K}$ | 2 * | mM |
| Parameter for the pump voltage dependence | $a$ | 0.39 * | |
| Parameter for the pump voltage dependence | $b$ | 1.28 * | |
| Extracellular K+ diffusion rate | $\varepsilon$ | $5.10^{-4}$ * | $ms^{-1}$ |
| K+ bath concentration | $K_{bath}$ | 3.5 * | mM |
| **Pyramidal neuron** | | | |
| Time constant for the decay of $S_e$ | $\tau_e$ | 3 * | ms |
| Voltage threshold defining for counting spikes | $V_{thres,e}$ | 20 # | mV |
| Fast inactivating Na+ maximal conductance | $g_{Na,FI,e}$ | 100 * | $mS.cm^{-2}$ |
| Delayed rectifier K+ maximal conductance | $g_{K,DR,e}$ | 80 * | $mS.cm^{-2}$ |
| Ca2+-activated K+ maximal conductance | $g_{K,AHP,e}$ | 1 * | $mS.cm^{-2}$ |
| Ca2+-activated K+ half activation $[Ca^{2+}]_e$ | $K_{Ca}$ | 0.001 * | mM |
| $Na^+$ leak conductance | $g_{Na,L,e}$ | 0.05 # | $mS.cm^{-2}$ |
| $K^+$ leak conductance | $g_{K,L,e}$ | 0.01 # | $mS.cm^{-2}$ |
| $Cl^-$ leak conductance | $g_{Cl,L,e}$ | 0.05 # | $mS.cm^{-2}$ |
| KCC2 cotransporter strength | $\rho_{KCC}$ | $6 \times 10^{-5}$ # | $mM.ms^{-1}$ |
| NKCC1 cotransporter strength | $\rho_{NKCC}$ | 10 # | $mM.ms^{-1}$ |
| NKCC1 cotransporter half activation $[K^+]$ | $K_{NKCC,K}$ | 16 * | mM |
| Glutamatergic current maximal conductance | $g_{GLU,e}$ | 0.1 * | $mS.cm^{-2}$ |
| GABAergic current maximal conductance | $g_{GABA,e}$ | 2.5 * | $mS.cm^{-2}$ |
| External glutamatergic conductance | $g_{D,e}$ | 0-0.3 * | $mS.cm^{-2}$ |
| $Ca^{2+}$ maximal conductance | $g_{Ca,e}$ | 1 * | $mS.cm^{-2}$ |
| $Ca^{2+}$ reversal potential | $E_{Ca,e}$ | 120 * | mV |
| Time constant for $Ca^{2+}$ extrusion and buffering | $\tau_{Ca}$ | 80 * | ms |

**Note**. Parameters marked with * were taken from Lemaire et al (2021), parameters marked with # symbol were adjusted to maintain system stability and preserve the resting membrane potential at -55.54 mV, the remaining unmarked parameters were taken from (Oruro et al. 2020) .

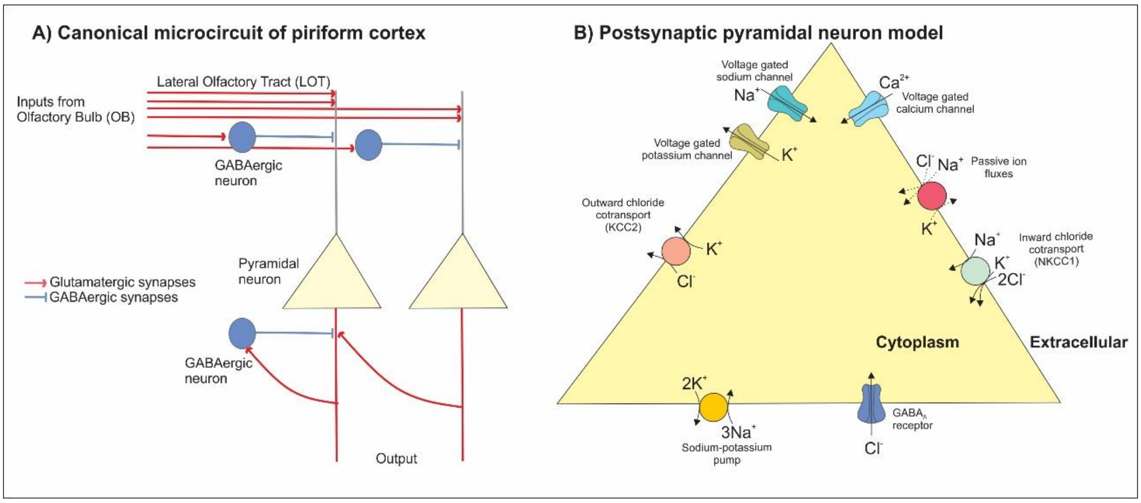

**Figure S1**. **Model representation. A)** Schematic representation of the canonical three-layered circuit of anterior piriform cortex (aPC). Pyramidal neurons are located in the deep layers and receive direct input from the olfactory bulb via the lateral olfactory tract (LOT), which projects to the superficial layers and establishes excitatory glutamatergic synapses. These LOT terminals also activate local GABAergic interneurons in the same layer, which in turn form GABAergic synapses onto the apical dendrites of pyramidal neurons. Additionally, pyramidal neurons form recurrent excitatory connections with one another in deeper layers and also synapse onto GABAergic interneurons that provide further inhibitory input (Oruro et al. 2020). **B**) Biophysical model of postsynaptic pyramidal neuron incorporating intracellular compartments connected by ion channels, transporters, and synaptic receptors. The model includes voltage-gated $Na^+$, $K^+$, and $Ca^{2+}$ channels, the $Na^+/K^+$ ATPase pump, chloride cotransporters (NKCC1 and KCC2), and GABAA receptor channels, allowing simulation of ion fluxes across the membrane. Based on Lemaire et al (2021).

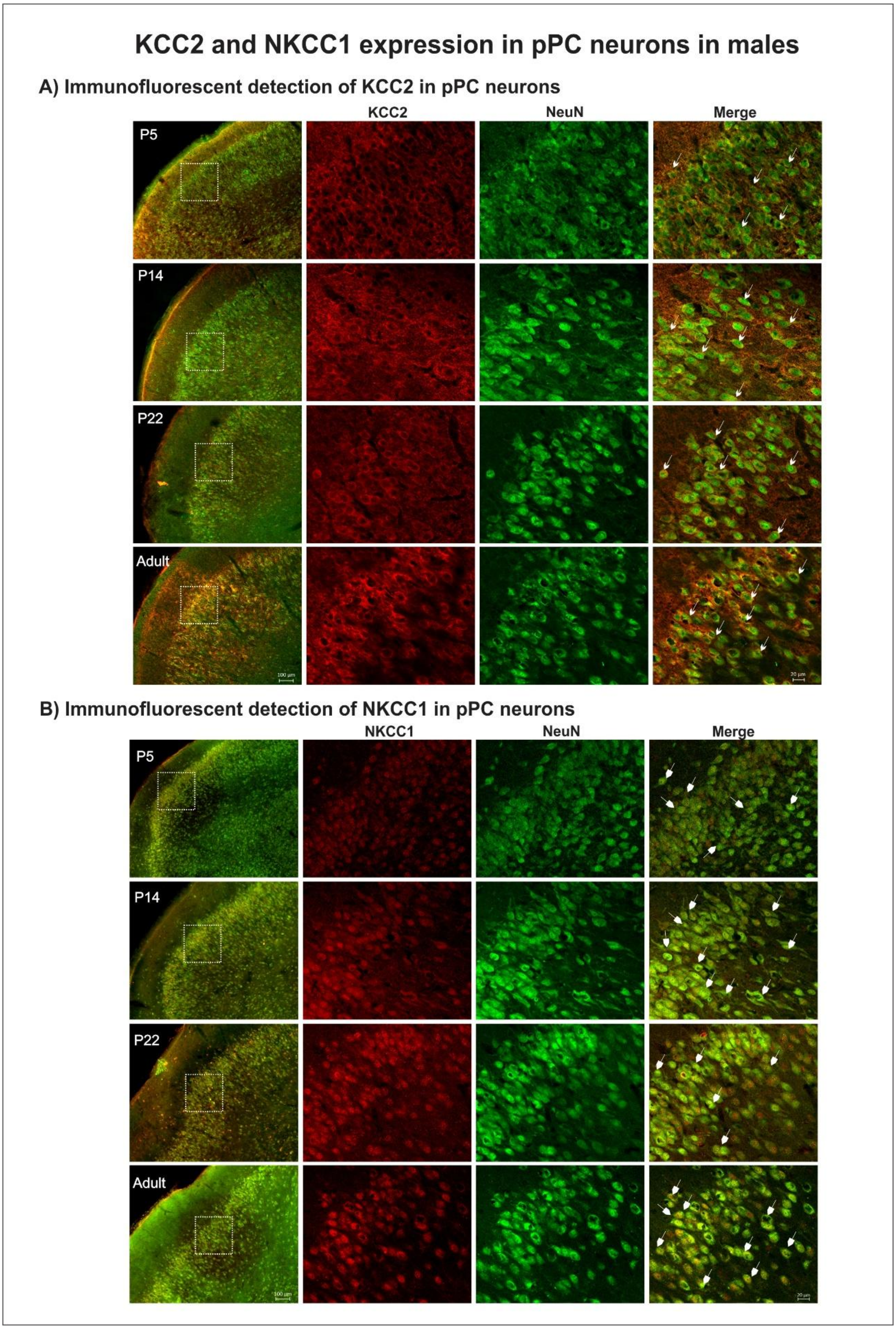


**Figure S2**. **Immunofluorescence images of KCC2 and NKCC1 expression in the posterior piriform cortex (pPC) neurons of male pups at P5, P14, P22, and adult**. **A**) Representative images showing KCC2 (red) and NeuN (green) immunoreactivity in the pPC at P5, P14, P22, and adult. **B**) Corresponding images showing NKCC1 (red) and

NeuN (green) expression in the same region and ages. For each age, overview panels (left column) show the location of the pPC region (scale bar = 100 µm). High-magnification images highlight immunolabeling in layers 2/3 (scale bar, 20 µm), which shows separate channels for KCC2 or NKCC1, NeuN, and the merged panel. White arrows in the merge panels indicate neurons co-expressing the chloride cotransporter and NeuN.

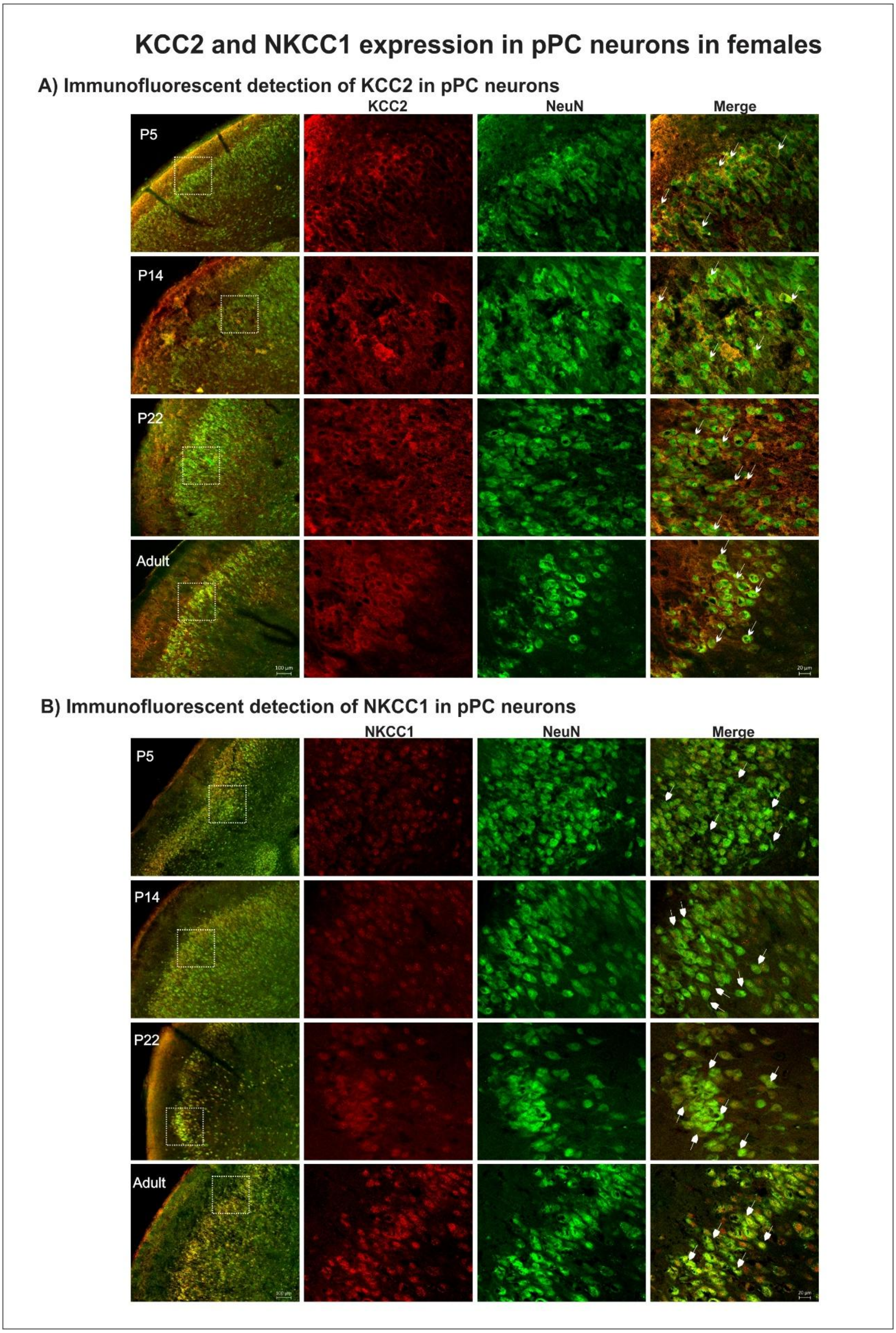


**Figure S3. Immunofluorescence images of KCC2 and NKCC1 expression in the posterior piriform cortex (pPC) neurons of female pups at P5, P14, P22, and adult.** **A**) Representative images showing KCC2 (red) and NeuN (green) immunoreactivity in the pPC at P5, P14, P22, and adult.**B**) Corresponding images showing NKCC1 (red) and NeuN (green) expression in the same region and ages. For each age, overview panels (left column) show the location of the pPC region (scale bar = 100 µm). High-magnification images highlight immunolabeling in layers 2/3 (scale bar, 20 µm), which shows separate channels for KCC2 or NKCC1, NeuN, and the merged panel. White arrows in the merge panels indicate neurons co-expressing the chloride cotransporter and NeuN.

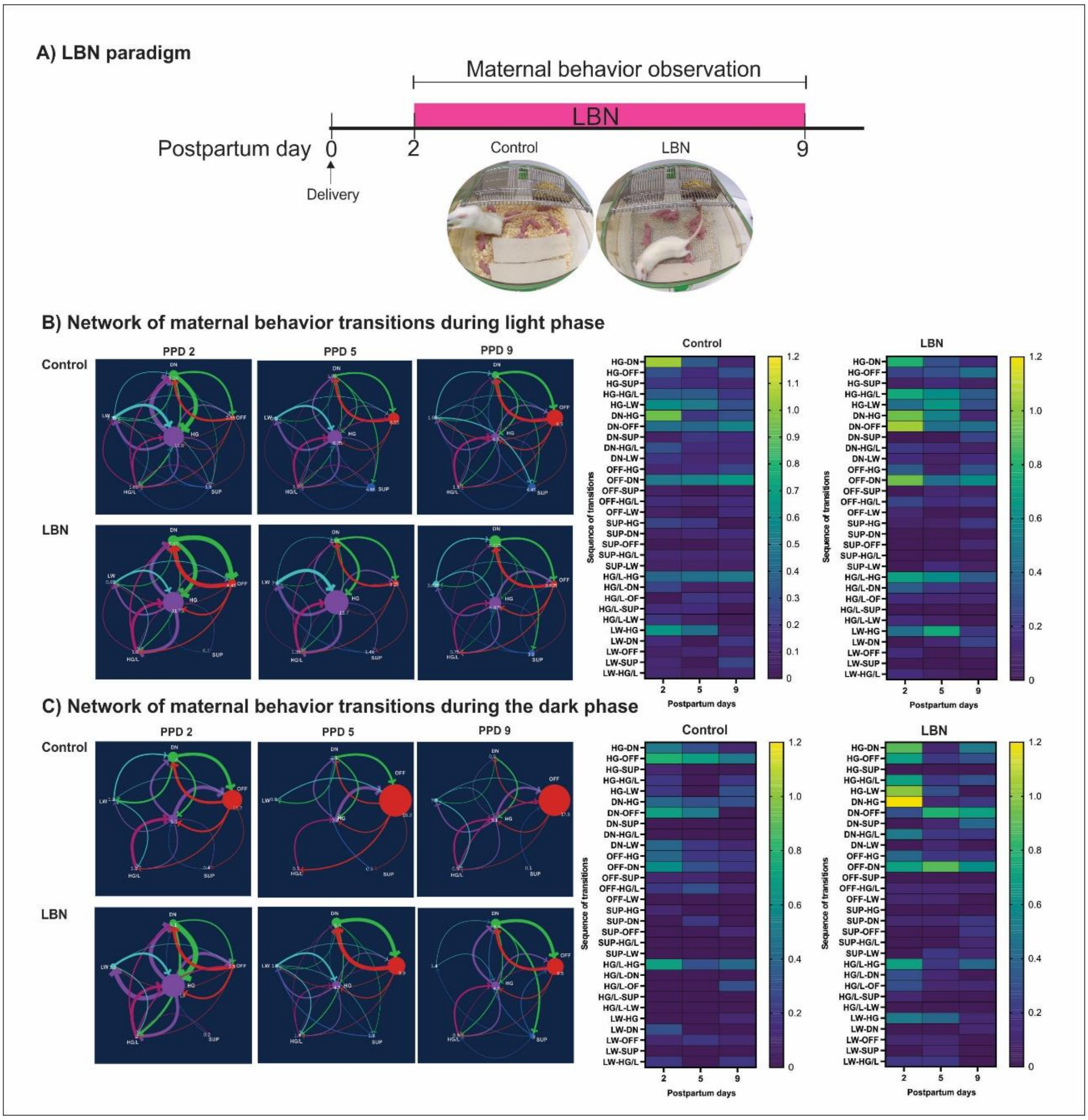


**Figure S4**. **Network transitions of maternal behavior across early postpartum days in control and LBN dams**. **A**) Experimental timeline of LBN exposure from postpartum day (PPD) 2 to 9. Representative networks of behavioral transitions during the light phase (**B**) and dark phase (**C**) on PPD 2, 5, and 9 in control and LBN conditions. Nodes represent averaged discrete maternal behaviors, color-coded as follows: HG, high crouch nursing posture (violet node); LW, low crouch nursing posture (cyan node); SUP, supine nursing posture (blue node); HG/L, high crouch nursing posture and licking pups at the same time (magenta node); DN, dam in nest without nursing pups (lime node); OFF, dam off the nest (red node). Arrows indicate the directions of transitions between behaviors, and arrow thickness indicates the frequency of each transition. LBN exposure increased transitions among maternal nursing postures. This is also represented in the adjacent heatmaps, where more intense yellow colors represent higher transition frequencies. These panels were constructed using data from Pardo et al. (2024)) and include a subset of time points from the original dataset. Adapted from **Figure 3** and **Figure 5** in (Pardo et al. 2024), an open-access publication under the terms of Creative Commons Attribution License.

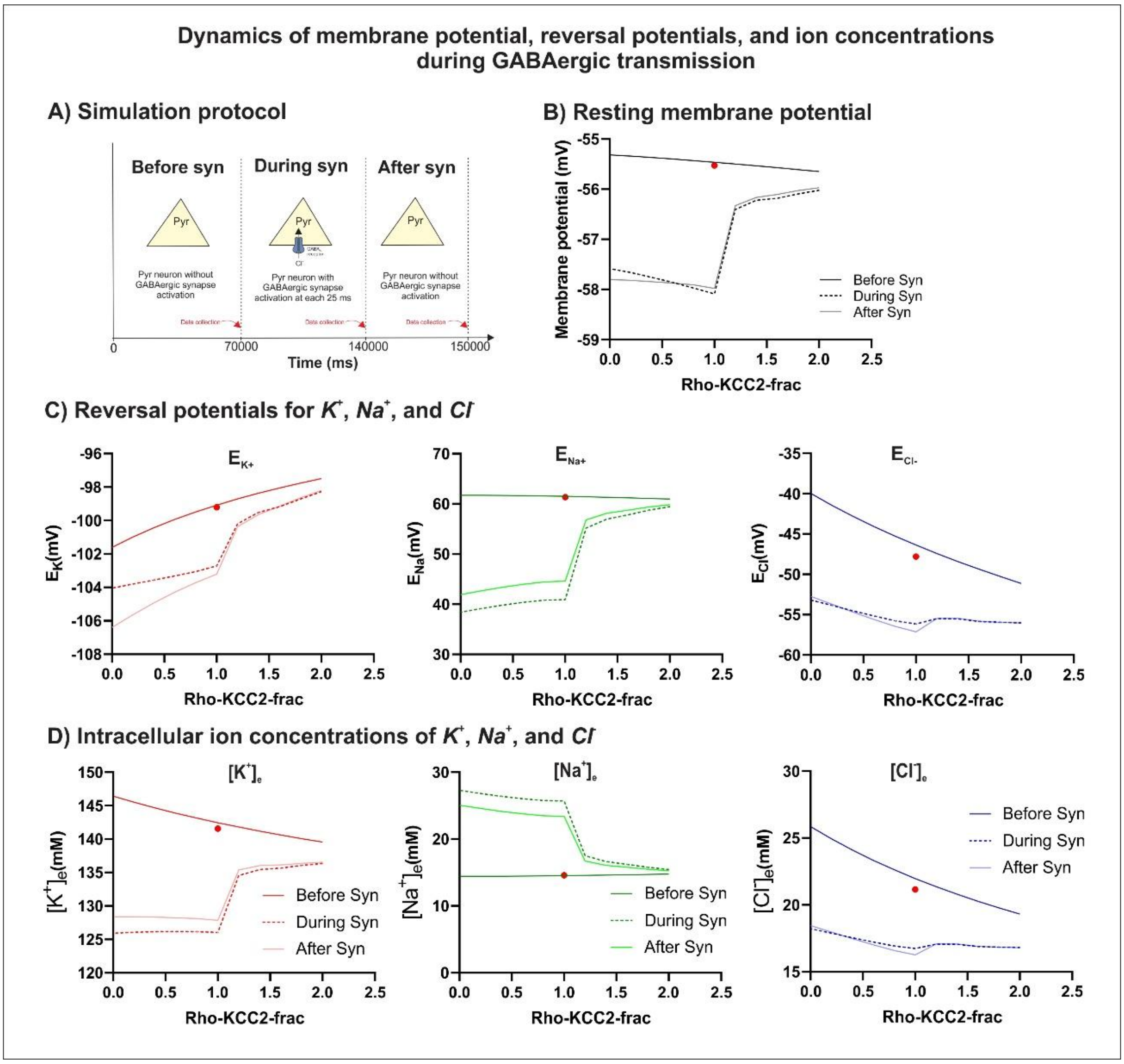


**Figure S5**. **Dynamic membrane potential and ion concentrations during GABAergic inputs. A**) Simulation protocol with three periods: before GABAergic synapse stimulation (*Before syn*, 0-70000 ms); during synapse stimulation (*During syn*, 70000-140000 ms), and after synapse stimulation (*After syn*, 14000-150000 ms). **B**) Variation of resting membrane potential as a function of KCC2 fraction measured before, during, and after GABAergic synapse stimulation. **C**) Reversal potential variation for potassium ($\boldsymbol{E_k}$), sodium ($\boldsymbol{E_{Na}}$), and chloride ($\boldsymbol{E_{Cl}}$) ions as a function. **D**) Intracellular ion concentration ([$K^+$], [$Na^+$], and [$Cl^-$]) as a function of KCC2, measured before, during, and after GABAergic synapse stimulation. For all experiments, the initial condition was set with KCC2 fraction equal to 1 and a membrane potential of -55.54 mV, corresponding to the experimental resting potential of aPC pyramidal neurons at P14-P17. This value is indicated as a red dot in each graph. Data points were collected at the end of each simulation period for the three conditions.

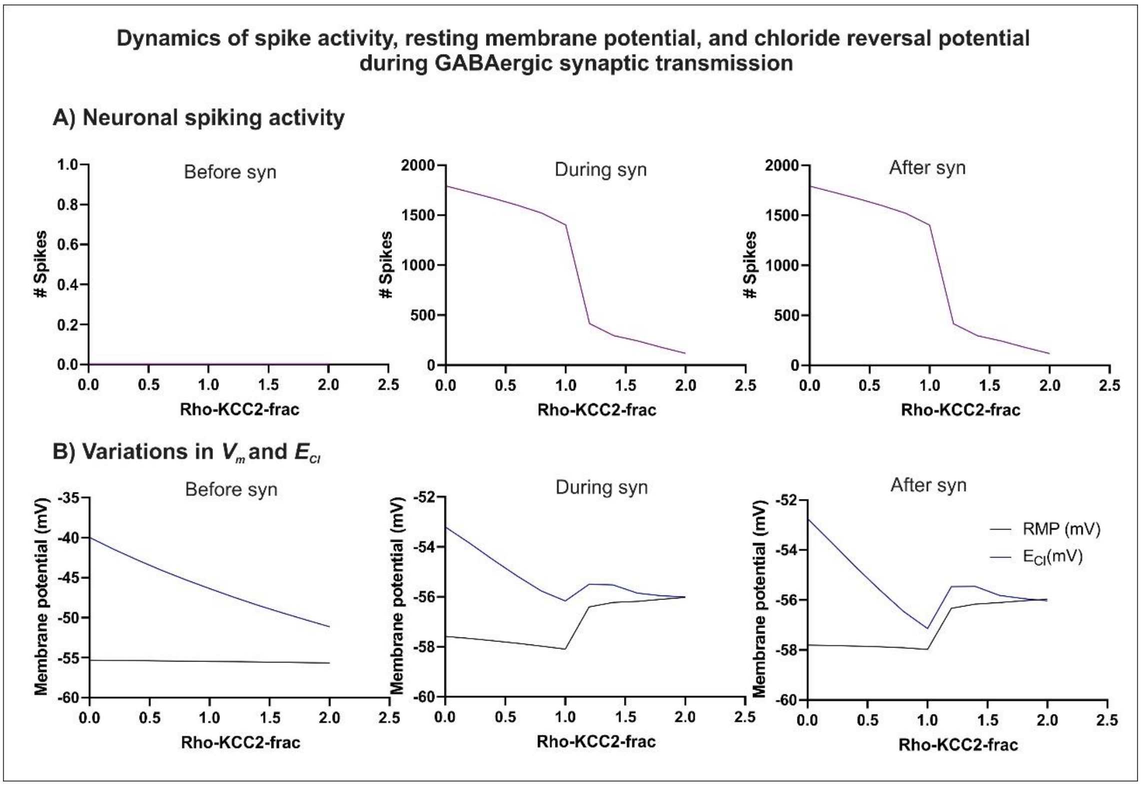


**Figure S6. Spiking activity, membrane potential, and chloride reversal potential during GABAergic inputs. A**) Spiking activity of the pyramidal neuron before GABAergic synapse stimulation (Before syn, 0-70000 ms); during synapse stimulation (During syn, 70000-140000 ms), and after synapse stimulation (After syn, 14000-150000 ms), as a function of KCC2 fraction. **B**) Variation of resting membrane potential (RPM) and chloride reversal potential (***$E_{Cl}$***) as a function of KCC2 fraction, measured before, during, and after GABAergic synapse stimulation. For all experiments, the initial condition was set with KCC2 fraction equal to 1 and a membrane potential of -55.54 mV, corresponding to the experimental resting potential of aPC pyramidal neurons at P14-P17. Data points were collected at the end of each simulation period for the three conditions.

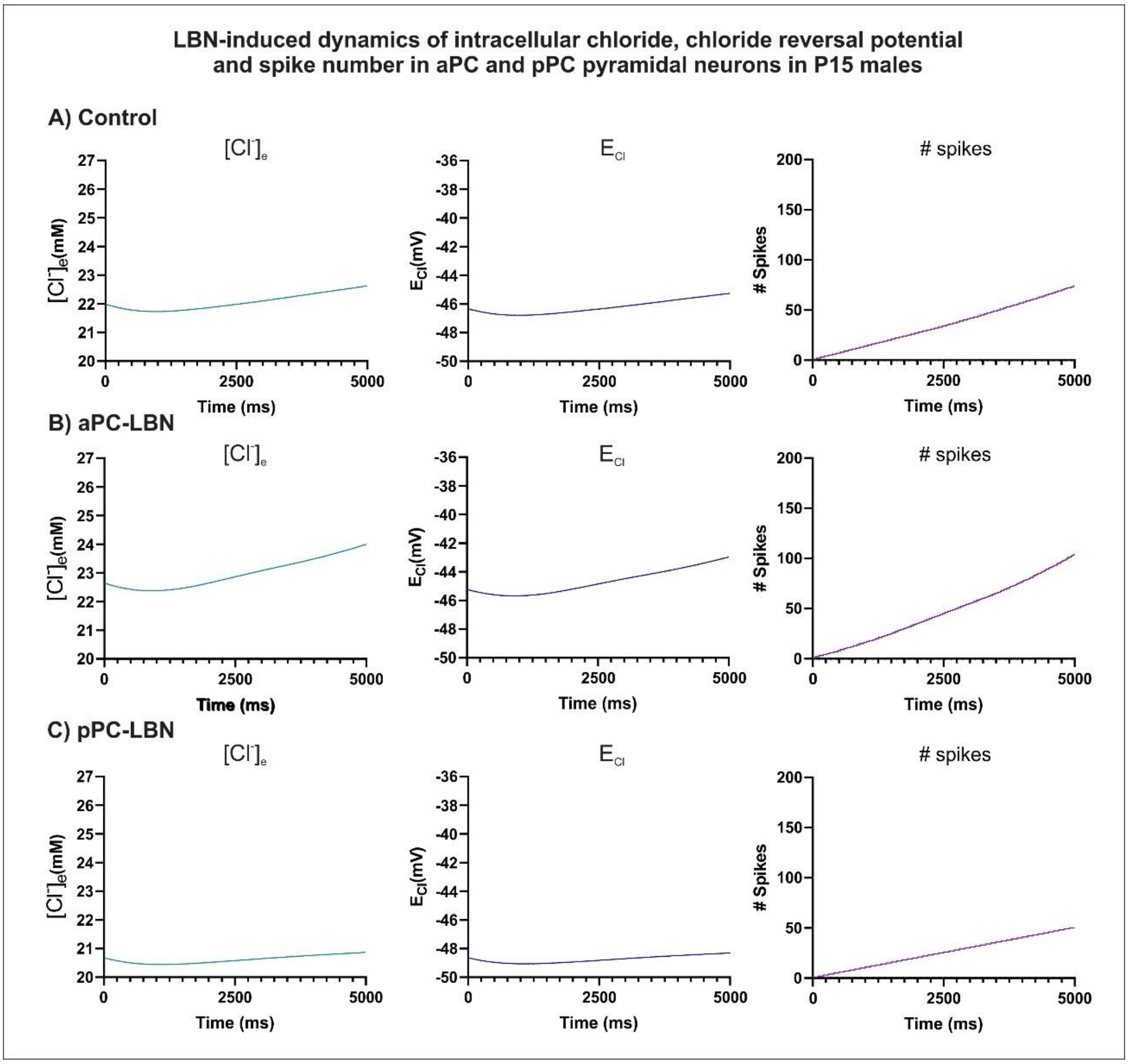


**Figure S7**. **Dynamics changes of intracellular ion concentration, reversal potential, and spiking of LBN-aPC and LBN-pPC neurons as a function of different fractions of KCC2/NKCC1 during GABAergic inputs in males. A**) Dynamics of intracellular chloride concentration, chloride reversal potential (***$E_{Cl}$***), and spike count of a control P15 aPC neuron with KKC2:NKCc1 fraction of 1:1 during GABAergic synapse simulation. **B**) Dynamics of intracellular chloride concentration, ***$E_{Cl}$***, and spike count of a P15 LBN-aPC neuron with KCC2:NKCC1 fraction of 0.80:0.59 during GABAergic synapse stimulation. **C**) Dynamics of intracellular chloride concentration, ***$E_{Cl}$***, and spike count of a P15 LBN-pPC neuron with KCC2:NKCC1 fraction of 1.45:0.90 during GABAergic synapse stimulation. For all experiments, simulations were preceded by a 70000 ms baseline period without GABAergic synapse stimulation, during which the KCC2 and NKCC1 fractions were set according to each experimental condition in male rats. The membrane potential was initialized at -55.54 mV, corresponding to the experimental resting potential of aPC pyramidal neurons at P14-P17. Data points were collected every 0.01 ms.

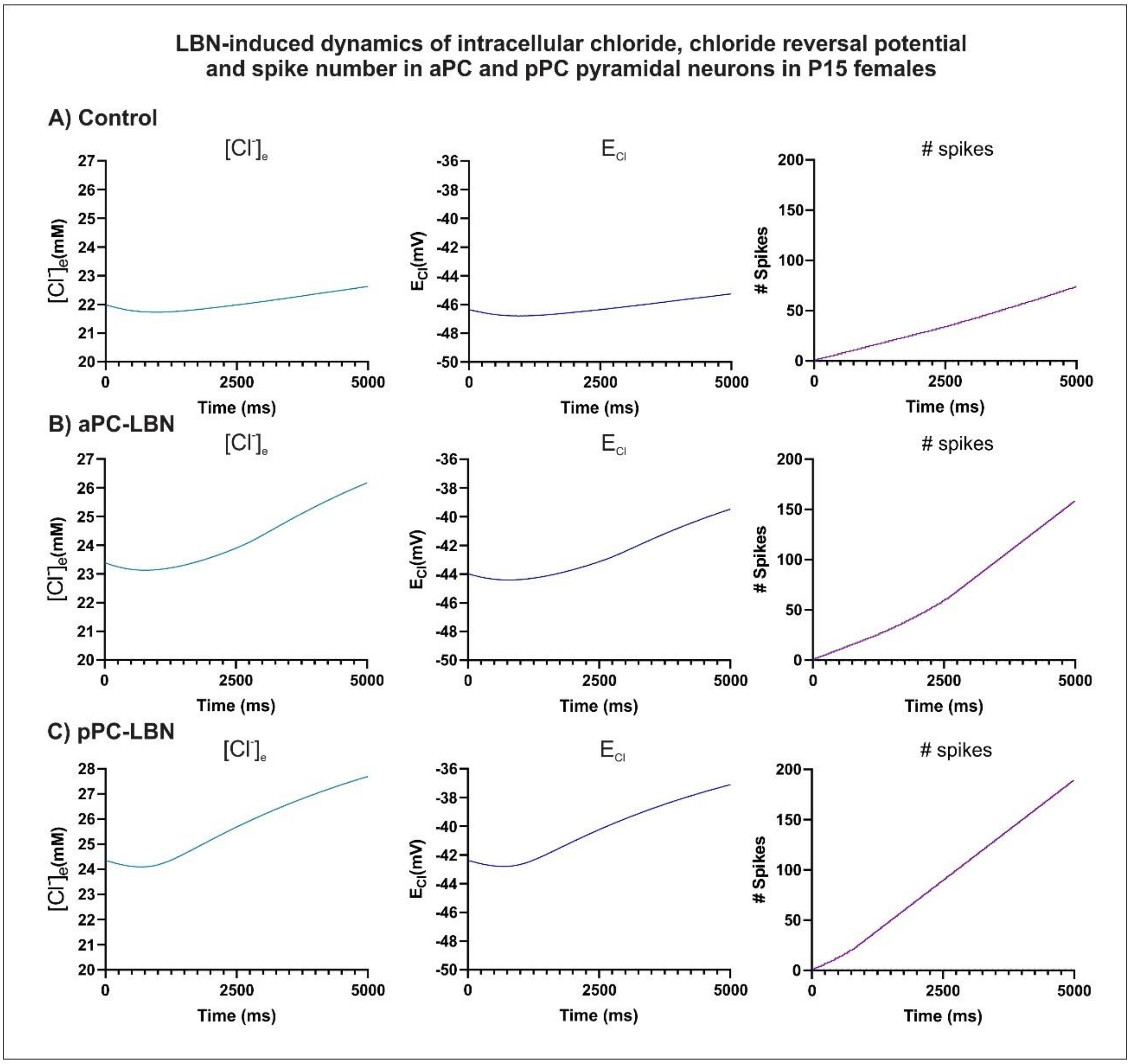


**Figure S8. Dynamics changes of intracellular ion concentration, reversal potential, and spiking of LBN-aPC and LBN-pPC neurons as a function of different fractions of KCC2/NKCC1 during GABAergic inputs in females. A**) Dynamics of intracellular chloride concentration, chloride reversal potential (***$E_{Cl}$***), and spike count of a control P15 aPC neuron with KKC2:NKCC1 fraction of 1:1 during GABAergic synapse simulation. **B**) Dynamics of intracellular chloride concentration, ***$E_{Cl}$,*** and spike count of a P15 LBN-aPC neuron with KCC2:NKCC1 fraction of 0.59:0.48 during GABAergic synapse stimulation. **C**) Dynamics of intracellular chloride concentration, ***$E_{Cl}$,*** and spike count of a P15 LBN-pPC neuron with KCC2:NKCC1 fraction of 0.34:0.88 during GABAergic synapse stimulation. For all experiments, simulations were preceded by a 70000 ms baseline period without GABAergic synapse stimulation, during which the KCC2 and NKCC1 fractions were set according to each experimental condition in female rats. The membrane potential was initialized at -55.54 mV, corresponding to the experimental resting potential of aPC pyramidal neurons at P14-P17. Data points were collected every 0.01 ms.
